# Virological characteristics of the novel SARS-CoV-2 Omicron variants including BA.2.12.1, BA.4 and BA.5

**DOI:** 10.1101/2022.05.26.493539

**Authors:** Izumi Kimura, Daichi Yamasoba, Tomokazu Tamura, Naganori Nao, Yoshitaka Oda, Shuya Mitoma, Jumpei Ito, Hesham Nasser, Jiri Zahradnik, Keiya Uriu, Shigeru Fujita, Yusuke Kosugi, Lei Wang, Masumi Tsuda, Mai Kishimoto, Hayato Ito, Rigel Suzuki, Ryo Shimizu, MST Monira Begum, Kumiko Yoshimatsu, Jiei Sasaki, Kaori Sasaki-Tabata, Yuki Yamamoto, Tetsuharu Nagamoto, Jun Kanamune, Kouji Kobiyama, Hiroyuki Asakura, Mami Nagashima, Kenji Sadamasu, Kazuhisa Yoshimura, Jin Kuramochi, Gideon Schreiber, Ken J Ishii, Takao Hashiguchi, The Genotype to Phenotype Japan (G2P-Japan) Consortium, Terumasa Ikeda, Akatsuki Saito, Takasuke Fukuhara, Shinya Tanaka, Keita Matsuno, Kei Sato

**Affiliations:** Division of Systems Virology, Department of Microbiology and Immunology, The Institute of Medical Science, The University of Tokyo, Tokyo, Japan; Faculty of Medicine, Kobe University, Kobe, Japan; Department of Microbiology and Immunology, Graduate School of Medicine, Hokkaido University, Sapporo, Japan; Division of International Research Promotion, International Institute for Zoonosis Control, Hokkaido University, Sapporo, Japan; One Health Research Center, Hokkaido University, Sapporo, Japan; Department of Cancer Pathology, Faculty of Medicine, Hokkaido University, Sapporo, Japan; Department of Veterinary Science, Faculty of Agriculture, University of Miyazaki, Miyazaki, Japan; Division of Molecular Virology and Genetics, Joint Research Center for Human Retrovirus infection, Kumamoto University, Kumamoto, Japan; Department of Clinical Pathology, Faculty of Medicine, Suez Canal University, Ismailia, Egypt; Department of Biomolecular Sciences, Weizmann Institute of Science, Rehovot, Israel; Graduate School of Medicine, The University of Tokyo, Tokyo, Japan; Institute for Chemical Reaction Design and Discovery (WPI-ICReDD), Hokkaido University, Sapporo, Japan; Division of Molecular Pathobiology, International Institute for Zoonosis Control, Hokkaido University, Sapporo, Japan; Institute for Genetic Medicine, Hokkaido University, Sapporo, Japan; Laboratory of Medical Virology, Institute for Life and Medical Sciences, Kyoto University, Kyoto, Japan; Department of Medicinal Sciences, Graduate School of Pharmaceutical Sciences, Kyushu University, Fukuoka, Japan; HiLung Inc., Kyoto, Japan; Division of Vaccine Science, Department of Microbiology and Immunology, The Institute of Medical Science, The University of Tokyo, Tokyo, Japan; International Vaccine Design Center, The Institute of Medical Science, The University of Tokyo, Tokyo, Japan; Tokyo Metropolitan Institute of Public Health, Tokyo, Japan; Interpark Kuramochi Clinic, Utsunomiya, Japan; Center for Animal Disease Control, University of Miyazaki, Miyazaki, Japan; Graduate School of Medicine and Veterinary Medicine, University of Miyazaki, Miyazaki, Japan; International Collaboration Unit, International Institute for Zoonosis Control, Hokkaido University, Sapporo, Japan; Division of Risk Analysis and Management, International Institute for Zoonosis Control, Hokkaido University, Sapporo, Japan; International Research Center for Infectious Diseases, The Institute of Medical Science, The University of Tokyo, Tokyo, Japan; CREST, Japan Science and Technology Agency, Kawaguchi, Japan

**Keywords:** SARS-CoV-2, COVID-19, Omicron, BA.4, BA.5, BA.2.12.1, BA.2, transmissibility, immune resistance, pathogenicity

## Abstract

After the global spread of SARS-CoV-2 Omicron BA.2 lineage, some BA.2-related variants that acquire mutations in the L452 residue of spike protein, such as BA.2.9.1 and BA.2.13 (L452M), BA.2.12.1 (L452Q), and BA.2.11, BA.4 and BA.5 (L452R), emerged in multiple countries. Our statistical analysis showed that the effective reproduction numbers of these L452R/M/Q-bearing BA.2-related Omicron variants are greater than that of the original BA.2. Neutralization experiments revealed that the immunity induced by BA.1 and BA.2 infections is less effective against BA.4/5. Cell culture experiments showed that BA.2.12.1 and BA.4/5 replicate more efficiently in human alveolar epithelial cells than BA.2, and particularly, BA.4/5 is more fusogenic than BA.2. Furthermore, infection experiments using hamsters indicated that BA.4/5 is more pathogenic than BA.2. Altogether, our multiscale investigations suggest that the risk of L452R/M/Q-bearing BA.2-related Omicron variants, particularly BA.4 and BA.5, to global health is potentially greater than that of original BA.2.

**Highlights:**

- Spike L452R/Q/M mutations increase the effective reproduction number of BA.2
- BA.4/5 is resistant to the immunity induced by BA.1 and BA.2 infections
- BA.2.12.1 and BA.4/5 more efficiently spread in human lung cells than BA.2
- BA.4/5 is more pathogenic than BA.2 in hamsters

## Introduction

Since the end of November 2021, the SARS-CoV-2 Omicron variant (B.1.1.529 and BA lineages) has spread worldwide and has outcompeted prior SARS-CoV-2 variants of concern (VOCs) such as Delta. After the surge of Omicron BA.1 variant, another Omicron variant, BA.2, outcompeted BA.1 and has become the most dominant variant in the world (Ito et al., 2022; UKHSA, 2022; Yamasoba et al., 2022a). Thereafter, as of May 2022, the Omicron subvariants that harbor the substitution at the L452 residue of spike (S) protein, such as BA.4 and BA.5, were frequently detected (Tegally et al., 2022; WHO, 2022). These observations suggest that these novel Omicron subvariants bearing mutations at the S L452 residue are more transmissible than Omicron BA.2. These recent developments have led the WHO to these Omicron subvariants bearing mutations at the S L452 residue, BA.4, BA.5, BA.2.12.1, BA.2.9.1 and BA.2.11, as VOC lineages under monitoring (VOC-LUM) on May 18, 2022 (WHO, 2022).

Resistance to antiviral humoral immunity can be mainly determined by the mutations in the S protein. For instance, Omicron BA.1 exhibits profound resistance to neutralizing antibodies induced by vaccination and natural SARS-CoV-2 infection as well as therapeutic monoclonal antibodies (Cao et al., 2021; Cele et al., 2021; Dejnirattisai et al., 2022; Garcia-Beltran et al., 2021; Liu et al., 2021; Meng et al., 2022; Planas et al., 2021; Takashita et al., 2022a; VanBlargan et al., 2022) and BA.2 (Bruel et al., 2022; Takashita et al., 2022b; Yamasoba *et al*., 2022a; Yamasoba et al., 2022b). In addition to immune evasion, the mutations in the S protein potentially modulate viral pathogenicity. Particularly, the fusogenicity of S protein in in vitro cell cultures is closely associated with viral pathogenicity in an experimental hamster model. For example, the Delta S is highly fusogenic in cell cultures and highly pathogenic in hamsters when compared to ancestral D614G-bearing B.1.1 S (Saito et al., 2022). In contrast, the Omicron BA.1 S is less fusogenic and pathogenic than B.1.1 S (Meng *et al*., 2022; Suzuki et al., 2022). Furthermore, we have recently demonstrated that the Omicron BA.2 S is more fusogenic and potentially confers the virus higher pathogenicity than Omicron BA.1 S (Yamasoba *et al*., 2022a).

Newly emerging SARS-CoV-2 variants need to be carefully and rapidly assessed for a potential increase in growth efficacy in the human population, pathogenicity and/or evasion from antiviral immunity. The substitution at the L452 residue of SARS-CoV-2 S protein was detected in Delta (L452R) and Lambda (L452Q) variants, which were previously classified as a VOC and a variant of interest (VOI), respectively (WHO, 2022). Importantly, we previously demonstrated that the L452R (Motozono et al., 2021) and L452Q (Kimura et al., 2022a) mutations increase viral infectivity by promoting the binding of S receptor binding domain (RBD) to human ACE2. We have recently characterized the virological features of SARS-CoV-2 Omicron BA.1 (Meng *et al*., 2022; Suzuki *et al*., 2022) and BA.2 (Yamasoba *et al*., 2022a). However, the impact of the substitution of S L452 residue on the virological characteristics of Omicron BA.2 remains unclear. Together with these findings, it is reasonable to assume that the novel BA.2-related Omicron variants bearing mutations at S L452 residue can be a potential risk for global health, and we herein elucidate the virological characteristics of these novel Omicron variants.

## Results

### Emergence of the BA.2-related Omicron bearing the L452R/Q/M mutations

Omicron substantially diversified during the epidemic. In South Africa, where Omicron was first reported at the end of November 2022 (NICD, 2021a; b), a variety of Omicron sublineages (BA.1–BA.5) continuously emerged (**Figures 1A and S1A**) (Tegally *et al*., 2022). Omicron BA.4 and BA.5 variants are closely related to each other and bear identical S protein (**Figure 1A**). Because BA.4 and BA.5 form a monophyletic clade with BA.2 (**Figure 1A**), we herein refer to BA.2, BA.4 and BA.5 as the BA.2-related Omicron variants. Compared to the BA.2 S, the BA.4/5 S harbors the L452R, HV69-70del, and F486V mutations as well as a revertant R493Q mutation. Notably, not only the BA.4 and BA.5 lineages, several BA.2 sublineages that bear the mutations at the S L452 residue also emerged (**Figure 1B and Table S1**). In-depth tracing of the emergence of BA.2 variants bearing mutations at the S L452 residue detected seven common ancestry groups of the BA.2 variants bearing L452R, L452Q or L452M mutation in the S protein (**Figures 1B, S1B, and S1C and Table S2)**. As of May 15, 2022, the PANGO lineage (https://cov-lineages.org) annotates four out of the seven BA.2 sublineages bearing mutations at the S L452 residue: BA.2.9.1 (+S:L452M) in Denmark, BA.2.11 (+S:L452R) in France, BA.2.12.1 (+S:L452Q/S704L) in the USA, and BA.2.13 in Belgium (+S:L452M) (**Figures 1B, 1C and S1B**), while the other three lineages are not annotated yet (**Figure 1B**). On May 18, 2022, the WHO classified these six these L452R/M/Q-bearing BA.2-related Omicron variants, which include BA.4, BA.5, BA.2.9.1, BA.2.11, BA.2.12.1, BA.2.13 and as VOC-LUM (WHO, 2022). Most importantly, these L452R/M/Q-bearing BA.2-related variants have higher effective reproduction numbers (R_e_) than the original BA.2 (**Figure 1D and Table S3**). Particularly, the R_e_ values of BA.4, BA.5, and BA.2.12.1 are 1.19-, 1.21-, and 1.13-fold higher than that of BA.2, respectively (**Figure 1D**), and these three variants begun outcompeting original BA.2 in several countries (**Figures 1E and S1D**). As of May 15, 2022, BA.4, BA.5, and BA.2.12.1 have been detected in 20, 19, and 36 countries, respectively (**Table S4**). Altogether, our data indicate that multiple BA.2-related Omicron variants bearing the mutations at the S L452 residue independently emerged in several countries, and further predict that these Omicron variants, particularly BA.4, BA.5 and BA.2.12.1, will spread worldwide and become the next predominant variants in the near future.

**Figure 1.**
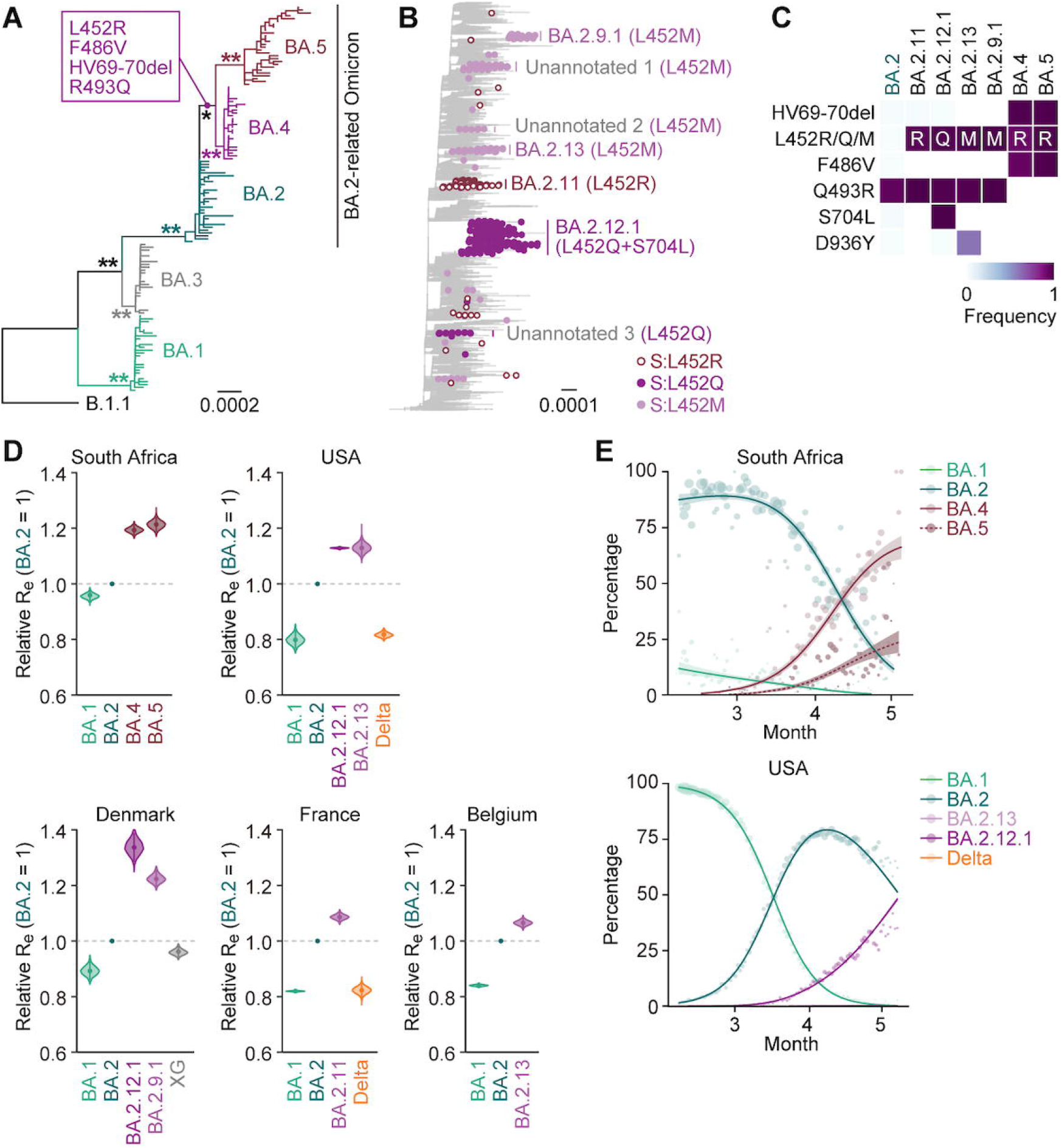
Epidemic of the BA.2-related Omicron variants bearing the S L452R/Q/M mutations. (**A**) A maximum likelihood (ML) tree of the Omicron lineages sampled from South Africa. The mutations acquired in the S proteins of BA.4 and BA.5 lineages are indicated in the panel. Note that R493Q is a reversion [i.e., back mutation from the BA.1–BA.3 lineages (R493) to the B.1.1 lineage (Q493)]. Bootstrap values, *, ≥ 0.85; **, ≥ 0.9. (**B**) An ML tree of BA.2. The BA.2 variants bearing mutations at the S L452 residue are indicated as colored dots, and the estimated common ancestry groups of the variants are indicated as vertical bars. The PANGO lineages are indicated in the panel. The mutations in the S proteins of each group are shown in parentheses. (**C**) Heatmap summarizing the frequency of amino acid substitutions. Substitutions detected in > 50% of sequences of any lineage are shown. (**D**) Estimated relative R_e_ of each viral lineage, assuming a fixed generation time of 2.1 days. The R_e_ value of BA.2 is set at 1. Posterior (violin), posterior mean (dot), and 95% confidential interval (CI) (line) are shown. (**E**) Epidemic dynamics of SARS-CoV-2 lineages. The results for up to five predominant lineages in South Africa (top) and the USA (bottom) are shown. The observed daily sequence frequency (dot) and the dynamics (posterior mean, line; 95% CI, ribbon) are shown. The dot size is proportional to the number of sequences. The BA.2 sublineages without mutations at the S L452 residue are summarized as “BA.2”. See also **Figure S1 and Tables S1–S4.**

### Immune resistance of the L452R/Q/M-bearing BA.2-related Omicron variants

We have recently demonstrated that BA.4/5 is more resistant to a therapeutic monoclonal antibody, cilgavimab, a component of Evusheld, than BA.2 (Yamasoba *et al*., 2022b). Additionally, Khan et al. have recently demonstrated that BA.4 and BA.5 are relatively resistant to the antiviral humoral immunity induced by BA.1 infection (Khan et al., 2022). To investigate the sensitivity of BA.2-related Omicron variants to antiviral humoral immunity, we prepared the pseudoviruses bearing the S proteins of these novel Omicron variants (BA.2.9.1, BA.2.11, BA.2.12.1 and BA.4/5) as well as their derivatives and original BA.2. Consistent with recent studies including ours (Khan *et al*., 2022; Yamasoba *et al*., 2022a), BA.2 was highly resistant to 14 convalescent sera from individuals who were infected with BA.1 (**Table S5**), and all BA.2-related Omicron variants tested were also resistant to these antisera (**Figure 2A**). In the case of the 16 sera infected with BA.1 from who were 2-dose or 3-dose vaccinated convalescents (i.e., BA.1 breakthrough infection) (**Table S5**), the sensitivity of BA.2.9.1 and BA.2.11 to these antisera was comparable to that of BA.2 (**Figure 2B**), and BA.2.12.1 was significantly (1.3-fold) more sensitive than BA.2 (**Figure 2B**; *P* = 0.021 by the Wilcoxon signed-rank test). In contrast, BA.4/5 was significantly (2.3-fold) more resistant to BA.1 breakthrough infection sera than BA.2 (**Figure 2B**; *P* < 0.0001 by the Wilcoxon signed-rank test), which is consistent with a recent study (Khan *et al*., 2022). The assay using the BA.2 S derivatives bearing individual mutations of BA.4/5 and the BA.1 breakthrough infection sera showed that the F486V mutation confers resistance, while the insertion of HV69-70del and R493Q mutations made the pseudovirus more sensitive (**Figure 2B**). Interestingly, the HV69-70del mutation is present in BA.1 but not in BA.2, while the R493Q is a reversive mutation in BA.4/5 (i.e., BA.1 and BA.2 are R493, while ancestral B.1.1 and BA.4/5 are Q493) (Yamasoba *et al*., 2022a). Therefore, these data suggest that the sensitivity of BA.2 HV69-70del mutant is attributed to BA.1 infection, while that of BA.2 R493Q is attributed to vaccination. Nevertheless, our data suggest that BA.4/5 is relatively more resistant to the immunity induced by BA.1 breakthrough infection than BA.2, and this resistance is attributed to the BA.4/5-specific F486V mutation.

**Figure 2.**
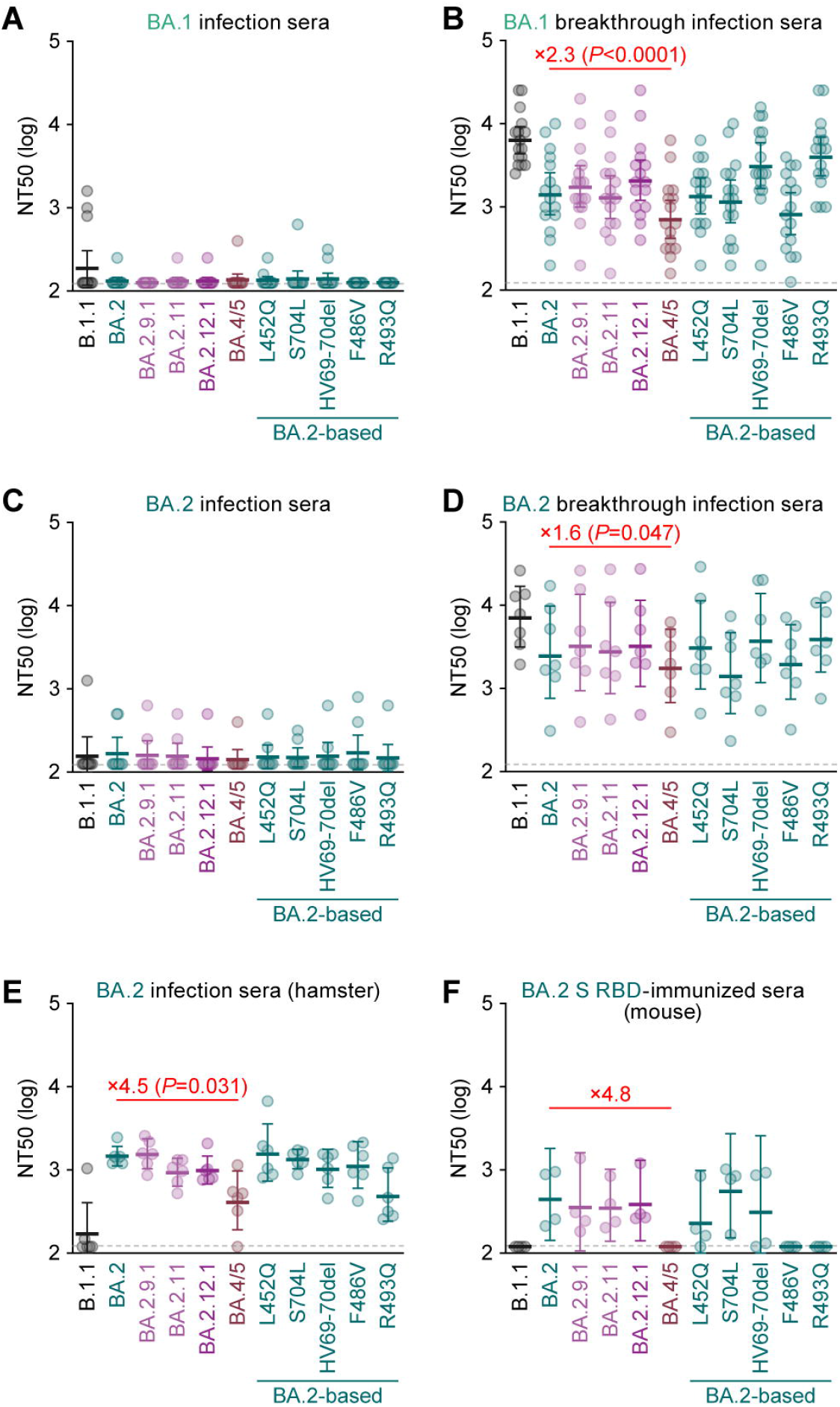
Immune resistance of the L452R/Q/M-bearing BA.2-related Omicron variants. Neutralization assays were performed with pseudoviruses harboring the S proteins of B.1.1 (the D614G-bearing ancestral virus), BA.1, BA.2 and the L452R/Q/M-bearing BA.2-related Omicron variants and the following sera. (**A and B**) Convalescent sera from individuals infected with BA.1 [14 not fully vaccinated donors (**A**) and 16 2/3-dose vaccinated donors (**B**)]. (**C and D**) Convalescent sera from individuals infected with BA.2 [9 not fully vaccinated donors (**C**) and 7 2/3-dose vaccinated donors (**D**)]. (**E**) Sera from BA.2-infected hamsters at 16 d.p.i. (6 hamsters). (**F**) Sera from mice immunized with BA.2 S RBD (4 mice). Assays with each serum sample were performed in triplicate to determine the 50% neutralization titer (NT50). Each dot represents one NT50 value, and the geometric mean and 95% CI are shown. The numbers indicate the fold changes of resistance versus each antigenic variant. The horizontal dashed line indicates the detection limit (120-fold). Statistically significant differences between BA.2 and other variants (*, *P* < 0.05) were determined by two-sided Wilcoxon signed-rank tests. Information on the vaccinated/convalescent donors is summarized in **Table S5**. See also **Table S5**.

We next tested the 16 sera infected with BA.2 from 8 convalescents who were not vaccinated and 1, 4 and 3 convalescents who were 1-dose, 2-dose, and 3-dose vaccinated convalescents, respectively (**Table S5**). As shown in **Figure 2C**, BA.2 convalescent sera did not exhibit antiviral effects against all variants tested including the D614G-bearing ancestral B.1.1. Although the BA.2 convalescent sera after breakthrough infection exhibited a stronger antiviral effect compared to the BA.2 convalescent sera without vaccination, BA.2 was 3.0-fold more resistant than B.1.1 (**Figure 2D**), suggesting that BA.2 infection does not induce efficient antiviral immunity. Nevertheless, BA.4/5 exhibited a significant (1.6-fold) resistance compared to BA.2 (**Figure 2D**; *P* = 0.047 by the Wilcoxon signed-rank test). To further address the possibility of evasion of BA.2-related Omicron variants from the immunity induced by the infection of original BA.2, we used the sera obtained from the hamsters infected with BA.2 (**Figure 2E**) (Yamasoba *et al*., 2022a) and the mice immunized with recombinant protein of BA.2 S receptor binding domain (RBD) (**Figure 2F**). In both cases, BA.4/5 evaded the BA.2-induced immunity (**Figures 2E and 2F**). Altogether, these results suggest that BA.4/5 is resistant to the immunity induced by BA.1 and BA.2.

### Virological features of the L452R/Q/M-bearing BA.2-related Omicron S

To investigate the virological characteristics of the L452R/Q/M-bearing BA.2-related Omicron variants, we measured pseudovirus infectivity using HOS-ACE2/TMPRSS2 cells (Kimura *et al*., 2022a; Motozono *et al*., 2021; Saito *et al*., 2022; Suzuki *et al*., 2022; Uriu et al., 2021; Yamasoba *et al*., 2022a). As shown in **Figure 3A**, all BA.2-related Omicron variants tested exhibited significantly higher infectivity than BA.2. The pseudovirus infectivity of BA.2.9.1, BA.2.11, BA.2.12.1 was comparable to that of ancestral D614G-bearing B.1.1, and notably, the infectivity of BA.4/5 pseudovirus was 18.3-fold higher than that of BA.2 pseudovirus (**Figure 3A**). The BA.2 derivatives bearing L452Q, HV69-70del and F486V mutations exhibited increased infectivity (**Figure 3A**). These results suggest that multiple mutations in the BA.4/5 S including HV69-70del, L452R and F486V increase pseudovirus infectivity. However, when we use HEK293-ACE2/TMPRSS2 cells and HEK293-ACE2 cells, which do not express endogenous TMPRSS2 on the cell surface (Yamasoba *et al*., 2022a), as target cells, the fold increase of pseudovirus infectivity of BA.2-related Omicron variants by the TMPRSS2 expression on the target cells was not observed (**Figure S2A**). These results suggest that the mutations detected in BA.2-related Omicron variants do not affect TMPRSS2 usage. Yeast surface display assay using SARS-CoV-2 S RBD and soluble human ACE2 (Dejnirattisai *et al*., 2022; Kimura *et al*., 2022a; Kimura et al., 2022b; Motozono *et al*., 2021; Yamasoba *et al*., 2022a; Zahradnik et al., 2021a) showed that the K_D_ value of BA.2 S RBD bearing L452R mutation is significantly lower than that of original BA.2 S RBD (**Figure 3B**), suggesting that the L452R mutation increases the binding affinity of BA.2 S RBD to human ACE2. On the other hand, the binding affinity of BA.4/5 S RBD to human ACE2 was comparable to that of BA.2 S RBD (**Figure 3B**).

**Figure 3.**
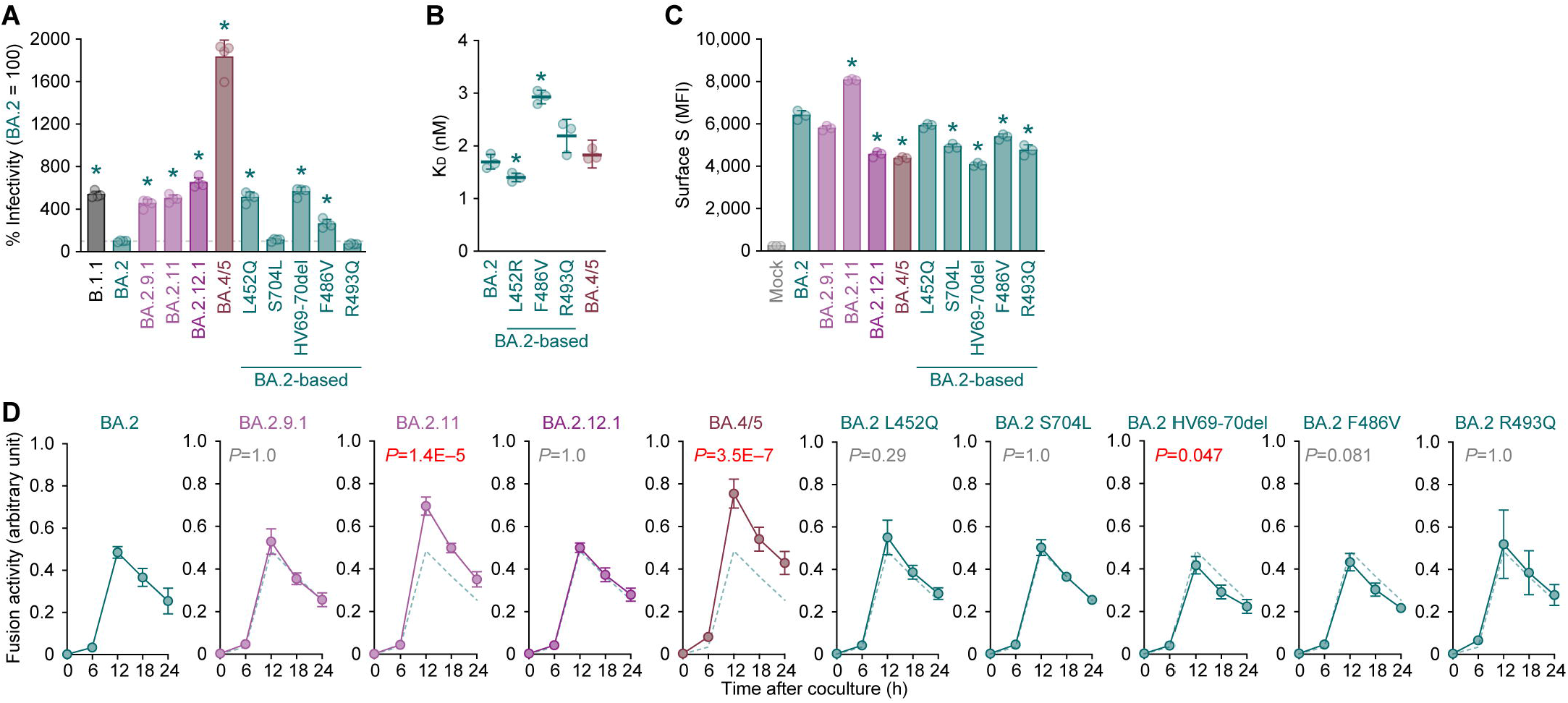
Virological features of the L452R/Q/M-bearing BA.2-related Omicron S. (**A**) Pseudovirus assay. The percent infectivity compared to that of the virus pseudotyped with BA.2 S are shown. (**B**) Binding affinity of SARS-CoV-2 S RBD to ACE2 by yeast surface display. The KD value indicating the binding affinity of the SARS-CoV-2 S RBD expressed on yeast binding to soluble ACE2 is shown. (**C and D**) S-based fusion assay. (**C**) S expression on the cell surface. Representative histograms stained with an anti-S1/S2 polyclonal antibody are shown in **Figure S2B**, and the summarized data are shown. In the left panel, the number in the histogram indicates mean fluorescence intensity (MFI). Gray histograms indicate isotype controls. (**D**) S-based fusion assay in Calu-3 cells. The recorded fusion activity (arbitrary units) is shown. The dashed green line indicates the results of BA.2. Assays were performed in quadruplicate (**A and D**) or triplicate (**B and C**), and the presented data are expressed as the average ± SD. Each dot indicates the result of an individual replicate. In **A–C**, statistically significant differences between BA.2 and other variants (*, *P* < 0.05 in **C**) were determined by two-sided Student’s *t* tests. In **D**, statistically significant differences between BA.2 and other variants across timepoints were determined by multiple regression. FWERs calculated using the Holm method are indicated in the figures. See also **Figure S2**.

We next analyzed the fusogenicity of BA.2-related Omicron variants by a cell-based fusion assay (Kimura *et al*., 2022b; Motozono *et al*., 2021; Saito *et al*., 2022; Suzuki *et al*., 2022; Yamasoba *et al*., 2022a). As shown in **Figures 3C and S2B**, the L452R mutation (specific in the BA.2.11 variant) significantly increased the S expression on the cell surface, while the L452Q and L452M (specific in the BA.2.9.1 variant) mutations did not. On the other hand, surface expression of the S proteins of BA.2.12.1 and BA.4/5 was significantly lower than that of original BA.2, and the decreased surface expression is attributed by the S704L mutation (specific in the BA.2.12.1 variant) and the HV69-70del, F486V and R493Q mutations (specific in the BA.4/5 variant) (**Figure 3C**). The cell-based fusion assay using Calu-3 cells as target cells showed that the fusogenicity of BA.2.11 S and BA.4/5 S was significantly greater than that of original BA.2 S, while the other mutations did not critically affect S-mediated fusogenicity (**Figure 3D**). When we use VeroE6/TMPRSS2 cells as target cells, all BA.2 derivatives bearing mutations at the L452 residue tested (i.e., L452R/M/Q mutations; BA.2.9.1, BA.2.11 and BA.2 L452Q) as well as the BA.4/5 significantly increased fusogenicity when compared to original BA.2, while the fusogenicity of the other mutants including BA.2.12.1 was comparable to original BA.2 (**Figure S2C**). Moreover, a coculture experiment using HEK293-ACE2/TMPRSS2 cells as the target cells (Suzuki *et al*., 2022; Yamasoba *et al*., 2022a) showed that the S proteins of BA.2.9.1, BA.2.11 and BA.4/5 but not BA.2.12.1 showed significantly increased fusogenicity than that of original BA.2 (**Figure S2D**). Altogether, these findings suggest that the L452R-bearing S proteins including BA.2.11 and BA.4/5 exhibited higher fusogenicity than BA.2 S in three independent experimental setups (**Figures 3D, S2C and S2D**).

### Growth capacity of the L452R/Q/M-bearing BA.2-related Omicron in vitro

We next prepared two sets of chimeric recombinant SARS-CoV-2 by reverse genetics (Kimura *et al*., 2022b; Motozono *et al*., 2021; Saito *et al*., 2022; Torii et al., 2021; Yamasoba *et al*., 2022a): one is based on an ancestral lineage A genome (strain WK-521; PANGO lineage A, GISAID ID: EPI_ISL_408667) and encodes *GFP* in the *ORF7a* frame, which was used in our previous studies (Kimura *et al*., 2022b; Saito *et al*., 2022; Yamasoba *et al*., 2022a) (**Figure 4A, left**), while the other is based on a clinical isolate of BA.2 (strain TY40-385; PANGO lineage BA.2, GISAID ID: EPI_ISL_9595859) (**Figure 4A, right**). The *S* genes of both recombinant viruses were swapped with those of BA.2-related Omicron variants: BA.2.11 (BA.2+L452R), BA.2.9.1 (BA.2+L452M), BA.2.12.1 (BA.2+L452Q/S704L) or BA.4/5 (BA.2+HV69-70del/L452R/F486V/R493Q) (**Figure 4A**). In the case of lineage A-based recombinant viruses (**Figure 4A, left**), the sizes of plaques formed by the infections of all BA.2-related Omicron variants (i.e., rBA.2.9.1 S-GFP, rBA.2.11 S-GFP, rBA.2.12.1 S-GFP and rBA.4/5 S-GFP) were significantly larger than that of rBA.2 S-GFP (**Figure 4B**). On the other hand, in the case of BA.2-based recombinant viruses (**Figure 4A, right**), the plaques formed by the infections of rBA.2.11 and rBA.4/5, which bear the L452R mutation, were larger than those formed by rBA.2 infection, and rBA.2.9.1 infection showed significantly smaller plaques than rBA.2 infection (**Figure 4C**). Corresponding to the results of the experiments using S expression plasmids (**Figures 3D, S2C and S2D**), these data suggest that the BA.4/5 S is more fusogenic than the BA.2 S.

**Figure 4.**
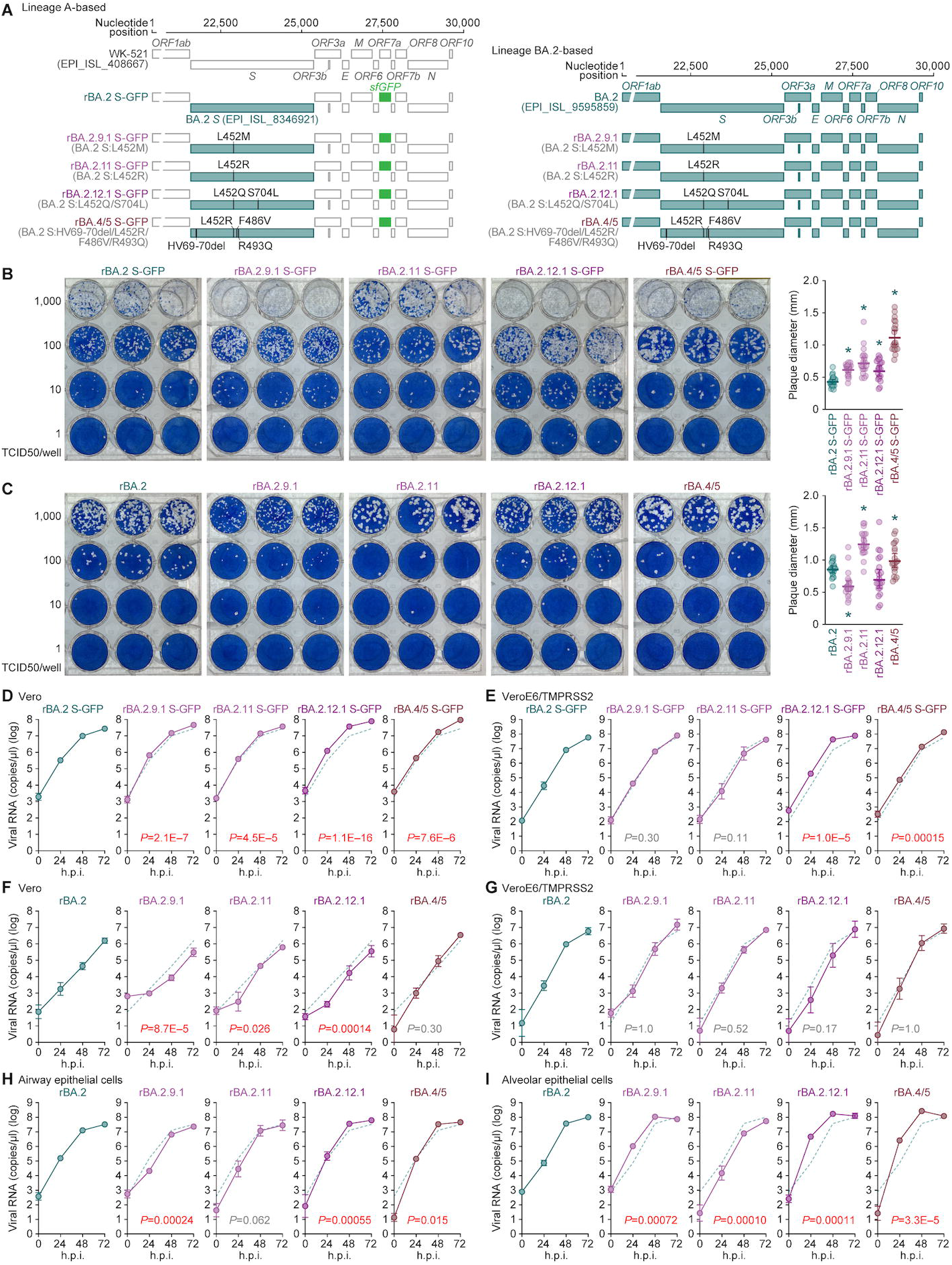
Growth capacity of the L452R/Q/M-bearing BA.2-related Omicron variants in vitro. (**A**) Scheme for the chimeric recombinant SARS-CoV-2 used in this study. The SARS-CoV-2 genome and its genes are shown. Left, the lineage A-based recombinant viruses. The template is SARS-CoV-2 strain WK-521 (PANGO lineage A, GISAID ID: EPI_ISL_408667), and the *S* genes were swapped with those of BA.2-related Omicron variants. *ORF7a* was swapped with the *sfGFP* gene. Right, the lineage BA.2-based recombinant viruses. The template is SARS-CoV-2 strain TY40-385 (PANGO lineage BA.2, GISAID ID: EPI_ISL_9595859), and the *S* genes were swapped with those of BA.2-related Omicron variants. The mutations based on the BA.2 S are summarized in parentheses. Note that the nucleotide sequences of two BA.2 isolates (GISAID IDs: EPI_ISL_8346921 and EPI_ISL_9595859) are identical. (**B and C**) Plaque assay. The lineage A-based (**B**) and lineage BA.2-based (**C**) recombinant viruses were respectively used, and VeroE6/TMPRSS2 cells were used for the target cells. Representative panels and summary of the recorded plaque diameters (20 plaques per virus) (most right) are shown. (**D-I**) Growth kinetics of chimeric recombinant SARS-CoV-2. Vero cells (**D and F**), VeroE6/TMPRSS2 cells (**E and G**), human iPSC-derived airway epithelial cells (**H**) and alveolar epithelial cells (**I**) were infected with the lineage A-based (**D and E**) and BA.2-based (**F-I**) chimeric recombinant SARS-CoV-2, and the copy numbers of viral RNA in the culture supernatant were routinely quantified by RT-qPCR. The dashed green line indicates the results of BA.2. In **B and C** (most right panels), each dot indicates the result of an individual plaque, and the presented data are expressed as the average ± SD. Statistically significant differences between BA.1 and BA.2 (*, *P* < 0.05) were determined by two-sided Mann–Whitney *U* tests. In **D-I**, assays were performed in quadruplicate, and the presented data are expressed as the average ± SD. Statistically significant differences between BA.2 and other variants across timepoints were determined by multiple regression. FWERs calculated using the Holm method are indicated in the figures.

To measure the growth kinetics of BA.2-related Omicron variants, the lineage A-based (**Figure 4A, left**) and lineage BA.2-based (**Figure 4A, right**) viruses were inoculated into the cells. As shown in **Figures 4D–4G**, the replication kinetics of L452R/M/Q-bearing BA.2-related Omicron variants were comparable to that of BA.2 regardless of their backbones. Notably, although the growth kinetics of BA.2-related Omicron variants, which are based on the BA.2 genome (**Figure 4A, right**), were relatively comparable in the culture of human airway epithelial cells derived from human induced pluripotent stem cells (iPSCs) (**Figure 4H**), rBA.2.9.1, rBA.2.12.1 and rBA.4/5 were significantly more efficiently replicated than rBA.2 in the human iPSC-derived alveolar epithelial cells (**Figure 4I**). Particularly, at 24 hours postinfection (h.p.i.), the levels of viral RNA in the supernatant of rBA.2.12.1- and rBA.4/5-infected cultures were 61-fold and 34-fold higher than that of rBA.2-infected culture (**Figure 4I**). These results suggest that rBA.2.12.1 and rBA.4/5 more efficiently replicate in human alveolar epithelial cells than BA.2.

### Virological features of BA.2.12.1 and BA.4/5 in vivo

To investigate the dynamics of viral replication of BA.2-related Omicron variants in vivo, we conducted hamster infection experiments using rBA.2, rBA.2.12.1 and rBA.4/5 (**Figure 4A, right**). Consistent with a recent report (Uraki et al., 2022), the rBA.2-infected hamsters did not exhibit apparent disorders based on the body weight, two surrogate markers of bronchoconstriction or airway obstruction [enhanced pause (Penh) and the ratio of time to peak expiratory follow relative to the total expiratory time (Rpef)], as well as decreased subcutaneous oxygen saturation (SpO_2_) (**Figure 5A**). Notably, the body weights of rBA.2.12.1-infected and rBA.4/5-infected hamsters were significantly lower than that of rBA.2-infected hamsters (**Figure 5A**). Additionally, the Rpef value of rBA.4/5-infected hamsters was significantly lower than that of rBA.2-infected hamsters (**Figure 5A**). These data suggest that the L452R/Q-bearing BA.2-related Omicron variants, particularly BA.4/5, exhibit higher pathogenicity than BA.2.

**Figure 5.**
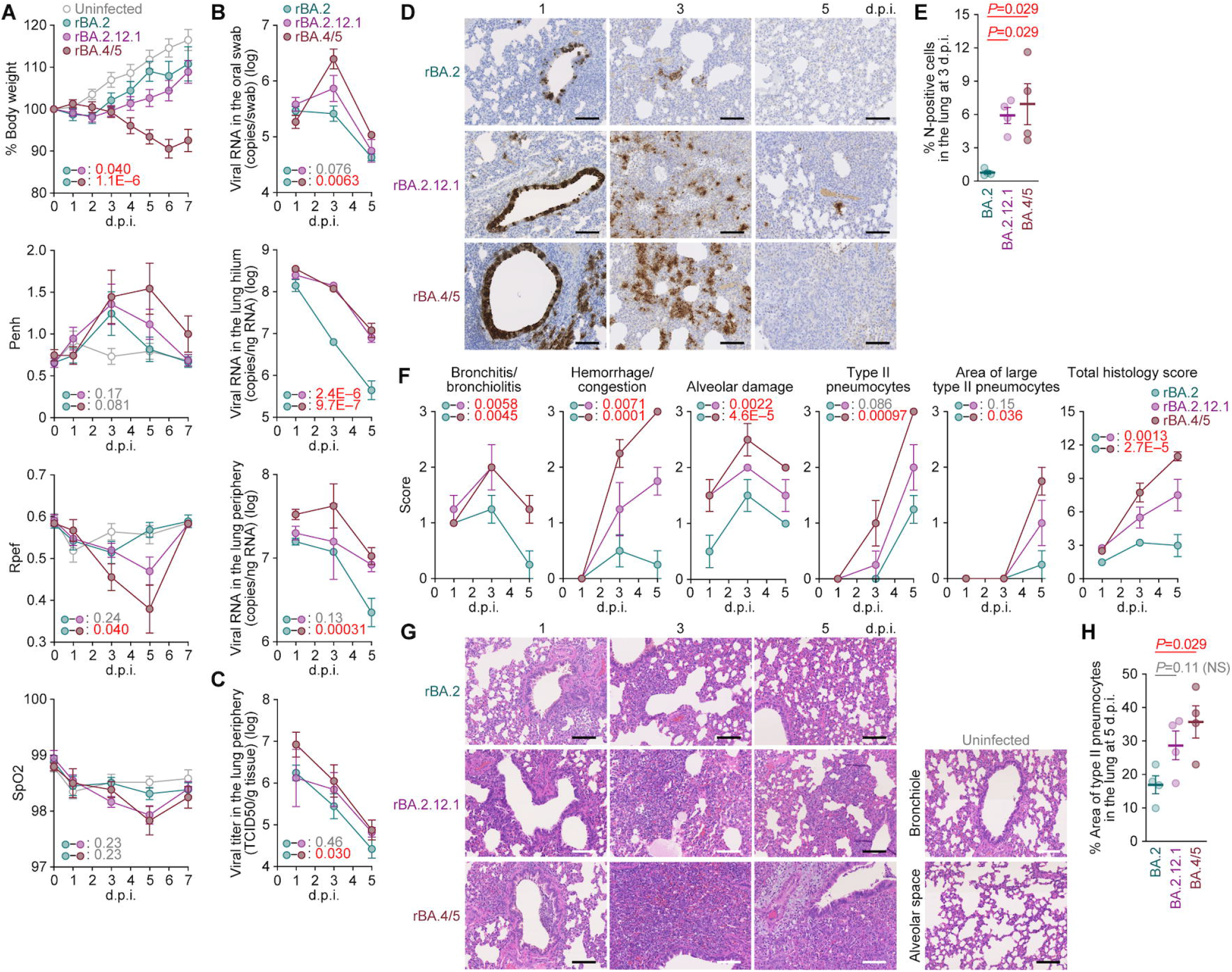
Virological features of BA.2.12.1 and BA.4/5 in vivo. Syrian hamsters were intranasally inoculated with rBA.2, rBA.2.12.1 and rBA.4/5 (summarized in Figure 4A**, right**). (**A**) Body weight, Penh, Rpef, and SpO_2_ values were routinely measured. Hamsters of the same age were intranasally inoculated with saline (uninfected). (**B**) Viral RNA loads in the oral swab (top), lung hilum (middle) and lung periphery (bottom). (**C**) Viral titers in the lung periphery. (**D**) IHC of the viral N protein in the lungs at 1, 3 and 5 d.p.i of all infected hamsters (n = 4 per viral strain). (**E**) Percentage of N-positive cells in whole lung lobes at 3 d.p.i. The raw data are shown in **Figure S3B**. (**F and G**) (**F**) Histopathological scoring of lung lesions. Representative pathological features are reported in our previous studies (Saito *et al*., 2022; Suzuki *et al*., 2022; Yamasoba *et al*., 2022a). (**G**) H&E staining of the lungs of infected hamsters. Uninfected lung alveolar space and bronchioles are also shown. (**H**) Type II pneumocytes in the lungs of infected hamsters. The percentage of the area of type II pneumocytes in the lung at 5 d.p.i. is shown. The raw data are shown in **Figure S3C**. Data are presented as the average (**A and B, top**, 6 hamsters per viral strain; **B, middle and bottom, C, E, F and H**, 4 hamsters per viral strain) ± SEM. In **E** and **H**, each dot indicates the result of an individual hamster. In **A–C and F**, statistically significant differences between rBA.2 and other variants across timepoints were determined by multiple regression. The 0 d.p.i. data were excluded from the analyses. FWERs calculated using the Holm method are indicated in the figures. In **E and H**, the statistically significant differences between rBA.2 and other variants were determined by a two-sided Mann–Whitney *U* test. In **D and G**, each panel shows a representative result from an individual infected hamster. Scale bars, 100 μm.\ See also **Figure S3**.

To analyze viral spread in the respiratory organs of infected hamsters, the viral RNA load and nucleocapsid (N) expression were assessed by RT-qPCR analysis of viral RNA and immunohistochemistry (IHC), respectively. As shown in **Figure 5B**, the viral RNA loads in the lung hilum of rBA.2.12.1- and rBA.4/5-infected hamsters were significantly higher than that of rBA.2-infected hamsters. Intriguingly, the viral RNA loads in the oral swab (**Figure 5B, top**) and lung periphery (**Figure 5B, bottom**) of rBA.4/5-infected hamsters were significantly higher than that of rBA.2, while those of rBA.2.12.1 were not significantly different from those of rBA.2. In particular, the levels of viral RNA in the lung periphery of rBA.4/5-infected hamsters at 3 and 5 days postinfection (d.p.i.) were 5.7-fold and 4.2-fold higher than those of rBA.2-infected hamsters, respectively (**Figure 5B, bottom**). The higher level of viral load in the lung periphery of rBA.4/5-infected hamsters than rBA.2-infected hamsters was also supported by the level of infectious viruses in these regions (**Figure 5C**). These results suggest that rBA.4/5 more efficiently spread in the lung of infected hamsters than rBA.2.

To address the possibility that rBA.4/5 spreads more efficiently than the BA.2, we investigated N protein positivity in the trachea and the lung area close to the hilum. At 1 d.p.i., there was no apparent difference in the N protein positivity in the tracheal epithelium among rBA.2-, rBA.2.12.1, and rBA.4/5-infected hamsters (**Figure S3A**). In the bronchial and bronchiolar epithelia, rBA.2.12.1 and BA.4/5 infections exhibited N-positive cells when compared to rBA.2 (**Figure 5D**). At 3 d.p.i., alveolar positivity was observed in rBA.2.12.1- and rBA.4/5-infected lungs but not in rBA.2-infected lungs (**Figure 5D**). Morphometry showed that the percentage of N-positive cells in rBA.2.12.1- and rBA.4/5-infected lungs was significantly higher than that in rBA.2-infected lungs at 3 d.p.i. (**Figures 5E and S3B**). At 5 d.p.i., N protein expression had almost disappeared in all infected lungs (**Figure 5D**). These data suggest that the rBA.2.12.1 and rBA.4/5 more efficiently spread in the lung tissues compared to rBA.2.

### Pathogenicity of BA.2.12.1 and BA.4/5 in vivo

To investigate the pathogenicity of BA.2.12.1 and BA.4/5, the right lungs of infected hamsters were collected at 1, 3, and 5 d.p.i. and subjected to histopathological analysis (**Figure 5F**) and hematoxylin and eosin (H&E) staining (**Figure 5G**) (Saito *et al*., 2022; Suzuki *et al*., 2022; Yamasoba *et al*., 2022a). Three histological parameters, including bronchitis/bronchiolitis, hemorrhage/congestion and alveolar damage, as well as the total score of histology of rBA.2.12-infected hamsters were significantly higher than those of rBA.2-infected hamsters (**Figures 5F and 5G**). More importantly, all histopathological parameters were highest in rBA.4/5-infected hamsters with statistical significances (**Figure 5F**). Furthermore, in the lungs of infected hamsters at 5 d.p.i., the level of inflammation with type II alveolar pneumocyte hyperplasia by rBA.4/5 infection was significantly higher than that by rBA.2 infection, while there was no statistically significant difference between rBA.2.12.1 and rBA.2 infections (**Figure 5H and S3C**). The relatively more severe disorders in the lungs of rBA.4/5-infected hamsters than those of rBA.2-infected hamsters (**Figures 5F–5H**) were supported by the more efficient spread of rBA.4/5 than rBA.2 in the infected lungs (**Figures 5B, bottom and 5C**). Altogether, these observations suggest that rBA.4/5 is highly pathogenic than rBA.2 in a hamster model.

## Discussion

Viral transmissibility, immune resistance and pathogenicity characterize the potential risk of new SARS-CoV-2 variant to global health. In this study, we investigated the virological characteristics of five “novel Omicron variants”, BA.2.9.1, BA.2.11, BA.2.12.1, BA.4 and BA.5. In these five variants, BA.4/5 renders highest potential risk in terms of the growth efficacy in the human population, resistance to antiviral humoral immunity, and pathogenicity in an experimental animal model.

As a common characteristic of the five BA.2-related Omicron variants focused in this study, the S proteins of these variants bear a mutation at the residue 452: BA.2.11 and BA.4/5, BA.2.9.1, and BA.2.12.1 respectively possess R, M and Q instead of parental L residue. The L452R and L452Q substitutions were present in previous a VOC and a VOI such as the Delta and Lambda variants (WHO, 2022), and in our previous studies (Kimura *et al*., 2022a; Motozono *et al*., 2021),we demonstrated that the L452R/Q mutation increases the binding affinity of S RBD to human ACE2, and thereby, increases pseudovirus infectivity. Here we demonstrated that the L452R mutation increases binding affinity to human ACE2 and pseudovirus infectivity even in the BA.2 S backbone. Therefore, together with our previous reports (Kimura *et al*., 2022a; Motozono *et al*., 2021), Additionally, the S proteins of BA.4 and BA.5 harbor the HV69-70del mutation, which was detected in the Alpha variant, a prior VOC. Consistent with a previous study (Meng et al., 2021), we demonstrated that the insertion of HV69-70del mutation increases pseudovirus infectivity. Altogether, multiple mutations in the S protein of BA.4/5 contribute to enhanced growth capacity in human lung cell culture and the lung of an experimental animal model.

In our previous studies that focused on Delta (Saito *et al*., 2022), Omicron BA.1 (Suzuki *et al*., 2022) and Omicron BA.2 (Yamasoba *et al*., 2022a), we suggested close association between viral fusogenicity in in vitro cell cultures and pathogenicity in vivo. For instance, a less fusogenic virus such as Omicron BA.1 was less pathogenic, while a more fusogenic virus such as Delta was more pathogenic (Saito *et al*., 2022; Suzuki *et al*., 2022). Here we demonstrated that the Omicron BA.4/5 variant is more fusogenic and pathogenic than the Omicron BA.2 variant. Consistent with previous findings (Saito *et al*., 2022; Suzuki *et al*., 2022; Yamasoba *et al*., 2022a), our data support the possibility that higher fusogenic virus tends to exhibit potentially higher pathogenicity at least in experimental animal models. Therefore, measuring the fusogenicity of viral S protein can be a rapid surrogate marker to assume potential viral pathogenicity.

A simplistic assumption without conclusive evidence implies that SARS-CoV-2 will evolve to attenuate its pathogenicity. However, we argue against this notion with at least three observations. First, the Delta variant exhibited relatively higher pathogenicity than the ancestral B.1 virus in an experimental animal model (Saito *et al*., 2022). Clinical studies also provide evidence suggesting the higher virulence of the Delta variant than other prior variants including the Alpha variant (Ong et al., 2021; Sheikh et al., 2021; Twohig et al., 2022). Second, although the Omicron BA.1 variant was less pathogenic than Delta and ancestral B.1.1 virus (Suzuki *et al*., 2022), the S protein of a subsequently spread variant, Omicron BA.2, acquired the potential to exhibit higher pathogenicity than that of Omicron BA.1 (Yamasoba *et al*., 2022a). Third, here we demonstrated that the Omicron BA.4/5 are more potentially pathogenic than Omicron BA.2. Therefore, our observations strongly suggest that SARS-CoV-2 does not necessarily evolve to attenuate its pathogenicity.

## Limitations of the study

In our previous study, we used a chimeric virus bearing the BA.2 *S* gene in a non-BA.2 (PANGO lineage A) genomic backbone and showed the BA.2 *S*-bearing chimeric virus is more pathogenic in infected hamsters than the BA.1 *S*-bearing chimeric virus (Yamasoba *et al*., 2022a). However, another study using a clinical isolate of BA.2 showed a comparable pathogenicity to a BA.1 clinical isolate (Uraki *et al*., 2022). This inconsistency of BA.2 pathogenicity found between our recent study (Yamasoba *et al*., 2022a) and other’s (Uraki *et al*., 2022) would be due to the difference of the viral sequence in the non-*S* region. In fact, there are 26 mutations in the non-*S* region between BA.2 and the non-BA.2 backbone (PANGO lineage A) used in our previous study (Yamasoba *et al*., 2022a) (**Table S1**). To avoid such inconsistency, here we used the recombinant viruses based on BA.2 for hamster experiments: compared to BA.2, the majority of BA.2.12.1 does not possess any mutations in the non-*S* region (**Table S1**), indicating that the BA.2-based recombinant virus encoding BA.2.12.1 S used for hamster experiments (rBA.2.12.1) is an authentic BA.2.12.1. Moreover, only six or two mutations were detected in the non-*S* regions of BA.4 and BA.5 genomes, respectively (**Table S1**). Therefore, it would be reasonable to assume that our findings in the use of recombinant viruses reflect the pathogenic potential of authentic BA.4/5 in a hamster model.

## STAR METHODS

- KEY RESOURCES TABLE
- RESOURCE AVAILABILITY

- Lead Contact
- Materials Availability
- Data and Code Availability
- EXPERIMENTAL MODEL AND SUBJECT DETAILS

- Ethics Statement
- Human serum collection
- Cell culture
- METHOD DETAILS

- Viral genome sequencing
- Phylogenetic and comparative genome analyses
- Definition of common ancestry groups of the BA.2 variants bearing mutations at position 452 in S
- Modeling the epidemic dynamics of SARS-CoV-2 lineages
- Plasmid construction
- Preparation of BA.2 S RBD
- Preparation of mouse sera
- Preparation of human airway and alveolar epithelial cells from human iPSCs
- Neutralization assay
- Pseudovirus infection
- Yeast surface display
- SARS-CoV-2 S-based fusion assay
- Coculture experiment
- SARS-CoV-2 reverse genetics
- SARS-CoV-2 preparation and titration
- Plaque assay
- SARS-CoV-2 infection
- RT-qPCR
- Animal experiments
- Lung function test
- IHC
- H&E staining
- Histopathological scoring
- QUANTIFICATION AND STATISTICAL ANALYSIS

## Supplemental Information

Additional Supplemental Items are available upon request.

## Author Contributions

Izumi Kimura, Daichi Yamasoba, Tomokazu Tamura, Shuya Mitoma, Hesham Nasser, Keiya Uriu, Shigeru Fujita, Yusuke Kosugi, Hayato Ito, Rigel Suzuki, Ryo Shimizu, MST Monira Begum, Terumasa Ikeda, Akatsuki Saito and Takasuke Fukuhara performed cell culture experiments.

Jiri Zahradnik and Gideon Schreiber performed a yeast surface display assay.

Jiei Sasaki, Kaori Sasaki-Tabata and Takao Hashiguchi prepared SARS-CoV-2 S RBD.

Kouji Kobiyama and Ken J Ishii prepared SARS-CoV-2 S RBD-immunized murine sera.

Daichi Yamasoba, Tomokazu Tamura, Naganori Nao, Mai Kishimoto, Rigel Suzuki, Kumiko Yoshimatsu and Keita Matsuno performed animal experiments. Yoshitaka Oda, Lei Wang, Masumi Tsuda and Shinya Tanaka performed histopathological analysis.

Hiroyuki Asakura, Mami Nagashima, Kenji Sadamasu, Kazuhisa Yoshimura performed viral genome sequencing analysis.

Yuki Yamamoto, Tetsuharu Nagamoto, and Jun Kanamune performed generation and provision of human iPSC-derived airway and alveolar epithelial cells.

Jin Kuramochi contributed clinical sample collection.

Jumpei Ito performed statistical, modelling, and bioinformatics analyses.

Jumpei Ito, Terumasa Ikeda, Akatsuki Saito, Takasuke Fukuhara, Shinya Tanaka, Keita Matsuno and Kei Sato designed the experiments and interpreted the results.

Jumpei Ito and Kei Sato wrote the original manuscript.

All authors reviewed and proofread the manuscript.

The Genotype to Phenotype Japan (G2P-Japan) Consortium contributed to the project administration.

## Acknowledgments

We would like to thank all members belonging to The Genotype to Phenotype Japan (G2P-Japan) Consortium. We thank Dr. Kenzo Tokunaga (National Institute for Infectious Diseases, Japan) and Dr. Jin Gohda (The University of Tokyo, Japan) and Dr. Hisashi Arase (Osaka University) for providing reagents. We also thank National Institute for Infectious Diseases, Japan, for providing a clinical isolate of Omicron BA.2 (GISAID ID: EPI_ISL_9595859). We gratefully acknowledge the laboratories responsible for obtaining the specimens and the laboratories where genetic sequence data were generated and shared via the GISAID Initiative. A full list of originating and submitting laboratories for the sequences used in our analysis can be found at https://www.gisaid.org using the EPI-SET-ID: EPI_SET_20220526dn. The super-computing resource was provided by Human Genome Center at The University of Tokyo.

This study was supported in part by AMED Research Program on Emerging and Re-emerging Infectious Diseases (JP21fk0108465, to Akatsuki Saito; JP20fk0108401, to Takasuke Fukuhara; JP20fk010847, to Takasuke Fukuhara; JP21fk0108617 to Takasuke Fukuhara; JP21fk0108146, to Kei Sato; JP22fk0108146, to Kei Sato; JP20fk0108413, to Terumasa Ikeda and Kei Sato; and JP20fk0108451, to G2P-Japan Consortium, Akatsuki Saito, Keita Matsuno, Takasuke Fukuhara, Terumasa Ikeda, and Kei Sato; JP20fk0108407, to Yuki Yamamoto and Tetsuharu Nagamoto); AMED Research Program on HIV/AIDS (JP21fk0410033, to Akatsuki Saito; JP22fk0410033, to Akatsuki Saito; JP22fk0410047, to Akatsuki Saito; JP22fk0410055, to Terumasa Ikeda; JP21fk0410039, to Kei Sato; and JP22fk0410039, to Kei Sato); AMED CRDF Global Grant (JP21jk0210039 to Akatsuki Saito; and JP22jk0210039 to Akatsuki Saito); AMED Japan Program for Infectious Diseases Research and Infrastructure (JP21wm0325009, to Akatsuki Saito; JP22wm0325009, to Akatsuki Saito; JP22wm0125008 to Keita Matsuno); JST A-STEP (JPMJTM20SL, to Terumasa Ikeda); JST SICORP (e-ASIA) (JPMJSC20U1, to Kei Sato); JST SICORP (JPMJSC21U5, to Kei Sato), JST CREST (JPMJCR20H4, to Kei Sato; JPMJCR20H8, to Takao Hashiguchi); JSPS KAKENHI Grant-in-Aid for Scientific Research C (19K06382, to Akatsuki Saito; 22K07103, to Terumasa Ikeda); JSPS KAKENHI Grant-in-Aid for Scientific Research B (21H02736, to Takasuke Fukuhara; 18H02662, to Kei Sato; and 21H02737, to Kei Sato); JSPS KAKENHI Grant-in-Aid for Early-Career Scientists (22K16375, to Hesham Nasser; 20K15767, Jumpei Ito); JSPS Fund for the Promotion of Joint International Research (Fostering Joint International Research) (18KK0447, to Kei Sato); JSPS Core-to-Core Program (A. Advanced Research Networks) (JPJSCCA20190008, to Kei Sato); JSPS Research Fellow DC1 (19J20488, to Izumi Kimura); JSPS Research Fellow DC2 (22J11578, to Keiya Uriu); JSPS Leading Initiative for Excellent Young Researchers (LEADER) (to Terumasa Ikeda); World-leading Innovative and Smart Education (WISE) Program 1801 from the Ministry of Education, Culture, Sports, Science and Technology (MEXT) (to Naganori Nao); Ministry of Education, Culture, Sports, Science and Technology grant 20H05773 (to Takao Hashiguchi); The Tokyo Biochemical Research Foundation (to Kei Sato); Mitsubishi Foundation (to Terumasa Ikeda); Shin-Nihon Foundation of Advanced Medical Research (to Terumasa Ikeda); Waksman Foundation of Japan (to Terumasa Ikeda); a Grant for Joint Research Projects of the Research Institute for Microbial Diseases, Osaka University (to Akatsuki Saito); an intramural grant from Kumamoto University COVID-19 Research Projects (AMABIE) (to Terumasa Ikeda); Intercontinental Research and Educational Platform Aiming for Eradication of HIV/AIDS (to Terumasa Ikeda).

## Consortia

The Genotype to Phenotype Japan (G2P-Japan) Consortium: Mai Suganami, Mika Chiba, Naoko Misawa, Ryo Yoshimura, Hirofumi Sawa, Kana Tsushima, Haruko Kubo, Zannatul Ferdous, Hiromi Mouri, Miki Iida, Keiko Kasahara, Koshiro Tabata, Mariko Ishizuka, Asako Shigeno, Isao Yoshida, So Nakagawa, Jiaqi Wu, Miyoko Takahashi, Kotaro Shirakawa, Akifumi Takaori-Kondo, Yasuhiro Kazuma, Ryosuke Nomura, Yoshihito Horisawa, Yusuke Tashiro, Yohei Yanagida, Yugo Kawai, Takashi Irie, Ryoko Kawabata, Otowa Takahashi, Kimiko Ichihara, Kazuko Kitazato, Haruyo Hasebe, Chihiro Motozono, Takamasa Ueno, Toong Seng Tan, Isaac Ngare, Erika P. Butlertanaka, Yuri L. Tanaka

---

**Table S1.** Summary of the mutations among Wuhan-Hu-1, original BA.2 and BA.2-related Omicron variants, related to Figure 1.

**Table S2.** The estimated common ancestry groups of BA.2 variants bearing the S L452 mutations, related to Figure 1.

**Table S3.** The estimated relative R_e_ values of viral lineages in each country, related to Figure 1.

**Table S4.** The detected sequence numbers of the BA.2-related Omicron lineages bearing the L452R/Q/M mutations in each country, related to Figure 1.

**Table S5.** Human sera used in this study, related to Figure 2.

**Table S6**. Primers used in this study, related to Figures 2–5.

**Table S7.** Summary of unexpected amino acid mutations detected in the working virus stocks, related to Figures 4–5.

**Figure S1. Phylogenetic analysis of the BA.2-related Omicron variants, related to Figure 1.**

(**A**) The mutation profile of Omicron lineages in South Africa, related to **Figure 1A**. Mutations detected in ≥5 sequences in the ML tree are summarized.

(**B**) The country and PANGO lineage of the BA.2 sequences in the ML tree, related to **Figure 1B**.

(**C**) Estimation of each common ancestry group of the S L452 mutation-bearing BA.2 variants. Amino acid at position 452 in S in each ancestral node was estimated by a Markov model, and the branches where the L452 mutation was acquired (red branches with asterisks) was estimated.

(**D**) Epidemic dynamics of SARS-CoV-2 lineages. The results for up to five predominant lineages in Denmark (left), France (middle) and Belgium (right) where the BA.2-related Omicron variants bearing the S L452R/Q/M mutations circulating are shown. The observed daily sequence frequency (dot) and the dynamics (posterior mean, line; 95% CI, ribbon) are shown. The dot size is proportional to the number of sequences. The BA.2 sublineages without mutations at the S L452 residue are summarized as “BA.2”.

**Figure S2. Virological features of BA.2 in vitro, related to Figure 3.**

(**A**) Fold increase in pseudovirus infectivity based on TMPRSS2 expression.

(**B**) S expression on the cell surface. Representative histograms stained with an anti-S1/S2 polyclonal antibody are shown. The number in the histogram indicates MFI. Gray histograms indicate isotype controls. The summarized data are shown in **Figure 3C**.

(**C**) S-based fusion assay in VeroE6/TMPRSS2 cells. The recorded fusion activity (arbitrary units) is shown. The dashed green line indicates the results of BA.2.

(**D**) Coculture of S-expressing cells with HEK293-ACE2/TMPRSS2 cells. Left, representative images of S-expressing cells cocultured with HEK293 cells (top) or HEK293-ACE2/TMPRSS2 cells (bottom). Nuclei were stained with Hoechst 33342 (blue). Right, size distribution of syncytia (green). The numbers in parentheses indicate the numbers of GFP-positive syncytia counted. Scale bars, 200 μm.

In **A and C**, assays were performed in quadruplicate, and the presented data are expressed as the average ± SD. In **A and D**, each dot indicates the result of an individual replicate.

In **C**, statistically significant differences between BA.2 and other variants across timepoints were determined by multiple regression. FWERs calculated using the Holm method are indicated in the figures.

In **D**, statistically significant differences between BA.2 and other variants (*, *P* < 0.05) were determined by two-sided Mann–Whitney *U* tests.

**Figure S3. Virological features of BA.2.12.1 and BA.4/5 in vivo, related to Figure 5.**

(**A**) IHC of the viral N protein in the middle portion of the tracheas of all infected hamsters (n = 4 per viral strain) at 1 d.p.i. Each panel shows a representative result from an individual infected hamster.

(**B**) Right lung lobes of hamsters infected with rBA.2, rBA.2.12.1 or rBA.4/5 (n = 4 per viral strain) at 3 d.p.i. were immunohistochemically stained with an anti-SARS-CoV-2 N monoclonal antibody. In each panel, IHC staining (top) and the digitalized N-positive area (bottom, indicated in red) are shown. The number in the bottom panel indicates the percentage of the N-positive area. Summarized data are shown in **Figure 5E**.

(**C**) Type II pneumocytes in the lungs of infected hamsters. Right lung lobes of hamsters infected with rBA.2, rBA.2.12.1 or rBA.4/5 (n = 4 per viral strain) at 5 d.p.i. In each panel, H&E staining (top) and the digitalized inflammation area (bottom, indicated in red) are shown. The number in the bottom panel indicates the percentage of the section represented by the indicated area (i.e., the area indicated in red within the total area of the lung lobe). Summarized data are shown in **Figure 5H**.

Scale bars, 1 mm.

## STAR METHODS

### KEY RESOURCES TABLE

### RESOURCE AVAILABILITY

#### Lead Contact

Further information and requests for resources and reagents should be directed to and will be fulfilled by the Lead Contact, Kei Sato.

#### Materials Availability

All unique reagents generated in this study are listed in the Key Resources Table and available from the Lead Contact with a completed Materials Transfer Agreement.

#### Data and Software Availability

The raw data of virus sequences analyzed in this study are deposited in the GitHub repository (https://github.com/TheSatoLab/BA.2_related_Omicrons). All databases/datasets used in this study are available from GISAID database (https://www.gisaid.org) and GenBank database (https://www.ncbi.nlm.nih.gov/genbank/). The accession numbers of viral sequences used in this study are listed in STAR METHODS.

The computational codes used in the present study are available in the GitHub repository (https://github.com/TheSatoLab/BA.2_related_Omicrons).

### EXPERIMENTAL MODEL AND SUBJECT DETAILS

#### Ethics statement

All experiments with hamsters were performed in accordance with the Science Council of Japan’s Guidelines for the Proper Conduct of Animal Experiments. The protocols were approved by the Institutional Animal Care and Use Committee of National University Corporation Hokkaido University (approval ID: 20-0123 and 20-0060). All experiments with mice were also performed in accordance with the Science Council of Japan’s Guidelines for the Proper Conduct of Animal Experiments. The protocols were approved by the Institutional Animal Experiment Committee of The Institute of Medical Science, The University of Tokyo (approval ID: PA21-39 and PA21-46). All protocols involving specimens from human subjects recruited at Kyoto University and Kuramochi Clinic Interpark were reviewed and approved by the Institutional Review Boards of Kyoto University (approval ID: G1309) and Kuramochi Clinic Interpark (approval ID: G2021-004). All human subjects provided written informed consent. All protocols for the use of human specimens were reviewed and approved by the Institutional Review Boards of The Institute of Medical Science, The University of Tokyo (approval IDs: 2021-1-0416 and 2021-18-0617), Kyoto University (approval ID: G0697), Kumamoto University (approval IDs: 2066 and 2074), and University of Miyazaki (approval ID: O-1021).

#### Human serum collection

Convalescent sera were collected from the following donors: vaccine-naïve individuals who had been infected with the Omicron BA.1 variant (n = 14; average age: 44, range: 16–73, 57% male), fully vaccinated individuals who had been infected with the Omicron BA.1 variant (n = 16; average age: 48, range: 20–76, 44% male), vaccine-naïve individuals who had been infected with the Omicron BA.2 variant (n = 9; average age: 34, range: 7–54, 44% male), and fully vaccinated individuals who had been infected with the Omicron BA.2 variant (n = 7; average age: 48, range: 29–71, 86% male). To identify the SARS-CoV-2 variants infecting patients, saliva was collected from COVID-19 patients during infection onset, and RNA was extracted using a QIAamp viral RNA mini kit (Qiagen, Cat# 52906) according to the manufacturer’s protocol. To detect the S E484A mutation (common in all Omicron variants including BA.1, BA.2 and the L452R/Q/M-bearing BA.2-related variants), a primer/probe E484A (SARS-CoV-2) (Takara, Cat# RC322A) was used. To detect the S R214EPE insertion (specific to BA.1, while undetectable in BA.2 and the L452R/Q/M-bearing BA.2-related variants), an in-house-developed protocol was used with the following primers and probe: Omi_ins214s-F1, 5’-TTC TAA GCA CAC GCC TAT TAT AGT GC-3’; Omi_ins214s-R1, 5’-TAA AGC CGA AAA ACC CTG AGG-3’; and Omi_ins214s, FAM-TGA GCC AGA AGA TC-MGB (Yamasoba *et al*., 2022a). To verify the absence of S L452R/Q/M mutation (specific to the L452R/Q/M-bearing BA.2-related variants, while undetectable in original BA.2), a L452R (SARS-CoV-2) primer/probe set v2 (Takara, Cat# RC346A) was used. Sera were inactivated at 56°C for 30 minutes and stored at –80°C until use. The details of the convalescent sera are summarized in **Table S5**.

#### Cell culture

HEK293T cells (a human embryonic kidney cell line; ATCC, CRL-3216), HEK293 cells (a human embryonic kidney cell line; ATCC, CRL-1573) and HOS-ACE2/TMPRSS2 cells (HOS cells stably expressing human ACE2 and TMPRSS2) (Ferreira et al., 2021; Ozono et al., 2021) were maintained in DMEM (high glucose) (Sigma-Aldrich, Cat# 6429-500ML) containing 10% fetal bovine serum (FBS, Sigma-Aldrich Cat# 172012-500ML), and 1% penicillin-streptomycin (PS) (Sigma-Aldrich, Cat# P4333-100ML). HEK293-ACE2 cells (HEK293 cells stably expressing human ACE2) (Motozono *et al*., 2021) was maintained in DMEM (high glucose) containing 10% FBS, 1 µg/ml puromycin (InvivoGen, Cat# ant-pr-1) and 1% PS. HEK293-ACE2/TMPRSS2 cells (HEK293 cells stably expressing human ACE2 and TMPRSS2) (Motozono *et al*., 2021) was maintained in DMEM (high glucose) containing 10% FBS, 1 µg/ml puromycin, 200 ng/ml hygromycin (Nacalai Tesque, Cat# 09287-84) and 1% PS. HEK293-C34 cells (*IFNAR1* KO HEK293 cells expressing human ACE2 and TMPRSS2 by doxycycline treatment) (Torii *et al*., 2021) were maintained in DMEM (high glucose) containing 10% FBS, 10 μg/ml blasticidin (InvivoGen, Cat# ant-bl-1) and 1% PS. Vero cells [an African green monkey (*Chlorocebus sabaeus*) kidney cell line; JCRB Cell Bank, JCRB0111] were maintained in Eagle’s minimum essential medium (EMEM) (Sigma-Aldrich, Cat# M4655-500ML) containing 10% FBS and 1% PS. VeroE6/TMPRSS2 cells (VeroE6 cells stably expressing human TMPRSS2; JCRB Cell Bank, JCRB1819) (Matsuyama et al., 2020) were maintained in DMEM (low glucose) (Wako, Cat# 041-29775) containing 10% FBS, G418 (1 mg/ml; Nacalai Tesque, Cat# G8168-10ML) and 1% PS. Calu-3 cells (a human lung epithelial cell line; ATCC, HTB-55) were maintained in EMEM (Sigma-Aldrich, Cat# M4655-500ML) containing 20% FBS and 1% PS. Calu-3/DSP_1-7_ cells (Calu-3 cells stably expressing DSP_1-7_) (Yamamoto et al., 2020) were maintained in EMEM (Wako, Cat# 056-08385) containing 20% FBS and 1% PS. 293S GnTI(-) cells (HEK293S cells lacking N-acetylglucosaminyltransferase (Kubota et al., 2016) were maintained in DMEM (Nacalai tesque, #08458-16 containing 2% FBS without PS. Expi293F cells (Thermo Fisher Scientific, Cat# A14527) were maintained in Expi293 expression medium (Thermo Fisher Scientific, Cat# A1435101). Human airway and alveolar epithelial cells derived from human induced pluripotent stem cells (iPSCs) were manufactured according to established protocols as described below (see “Preparation of human airway and alveolar epithelial cells from human iPSCs” section) and provided by HiLung Inc.

### METHOD DETAILS

#### Viral genome sequencing

Viral genome sequencing was performed as previously described (Meng *et al*., 2022; Motozono *et al*., 2021; Saito *et al*., 2022; Suzuki *et al*., 2022; Yamasoba *et al*., 2022a). Briefly, the virus sequences were verified by viral RNA-sequencing analysis. Viral RNA was extracted using a QIAamp viral RNA mini kit (Qiagen, Cat# 52906). The sequencing library employed for total RNA sequencing was prepared using the NEB next ultra RNA library prep kit for Illumina (New England Biolabs, Cat# E7530). Paired-end 76-bp sequencing was performed using a MiSeq system (Illumina) with MiSeq reagent kit v3 (Illumina, Cat# MS-102-3001). Sequencing reads were trimmed using fastp v0.21.0 (Chen et al., 2018) and subsequently mapped to the viral genome sequences of a lineage A isolate (strain WK-521; GISAID ID: EPI_ISL_408667) (Matsuyama *et al*., 2020) using BWA-MEM v0.7.17 (Li and Durbin, 2009). Variant calling, filtering, and annotation were performed using SAMtools v1.9 (Li et al., 2009) and snpEff v5.0e (Cingolani et al., 2012).

#### Phylogenetic and comparative genome analyses

To construct an ML tree of Omicron lineages (BA.1–BA.5) sampled from South Africa (shown in **Figure 1A**), the genome sequence data of SARS-CoV-2 and its metadata were downloaded from the GISAID database (https://www.gisaid.org/) (Khare et al., 2021) on April 23, 2022. We excluded the data of viral strains with the following features from the analysis: i) a lack collection date information; ii) sampling from animals other than humans, iii) >2% undetermined nucleotide characters, or iv) sampling by quarantine. From each viral lineage, 30 sequences were randomly sampled and used for tree construction, in addition to an outgroup sequence, EPI_ISL_466615, representing the oldest isolate of B.1.1 obtained in the UK. The viral genome sequences were mapped to the reference sequence of Wuhan-Hu-1 (GenBank accession number: NC_045512.2) using Minimap2 v2.17 (Li, 2018) and subsequently converted to a multiple sequence alignment according to the GISAID phylogenetic analysis pipeline (https://github.com/roblanf/sarscov2phylo). The alignment sites corresponding to the 1–265 and 29674–29903 positions in the reference genome were masked (i.e., converted to NNN). Alignment sites at which >50% of sequences contained a gap or undetermined/ambiguous nucleotide were trimmed using trimAl v1.2 (Capella-Gutierrez et al., 2009). Phylogenetic tree construction was performed via a three-step protocol: i) the first tree was constructed; ii) tips with longer external branches (Z score > 4) were removed from the dataset; iii) and the final tree was constructed. Tree reconstruction was performed by RAxML v8.2.12 (Stamatakis, 2014) under the GTRCAT substitution model. The node support value was calculated by 100 times bootstrap analysis.

To classify the BA.2 variants bearing mutations at the S L452 residue, we constructed an ML tree of BA.2 variants including those bearing mutations at the S L452 residue (shown in **Figure 1B**). For quality control, the BA.2 sequences without the S:N501Y and S:E484A mutations, characteristic mutations of Omicron, were removed from the dataset. Also, the BA.2 sequences with the S:HV69-70del mutation, a deletion mutation that is not present in BA.2 but in other Omicron lineages, were removed. To make a subset of BA.2 sequences representing the diversity of BA.2 for tree construction, we defined the “common amino acid haplotype” of BA.2 as described below. We first extracted the BA.2 sequences bearing mutations at position 452 in S. In these BA.2 variants, amino acid mutations (including substitutions, insertions, and deletions) present > 1% sequences were detected and referred to as the “common amino acid mutations”. According to the profile of the common amino acid mutations, a common amino acid haplotype, a set of common amino acid mutations present in each sequence, was determined for all BA.2 sequences. Finally, up to 20 sequences were randomly sampled from each unique common amino acid haplotype. As outgroup sequences, the oldest isolate of B.1.1 obtained in the UK (EPI_ISL_466615) and the oldest five BA.1 and BA.3 sequences sampled from South Africa after December 1, 2022, were used. The ML tree was constructed by the procedure described above. In the final set, 8,029 BA.2 sequences were included. Outgroup sequences are not displayed in **Figure 1B**.

#### Definition of common ancestry groups of the BA.2 variants bearing mutations at position 452 in S

According to the phylogenetic tree of BA.2 shown in **Figure 1B**, we defined common ancestry groups of the BA.2 variants bearing mutations at position 452 in S as the follow procedures. First, the ancestral state of the amino acid at position 452 in S at each node was estimated using a fixed-rates continuous-time Markov model (Mk model) implemented in the R package “castor” (**Figure S1C**) (Louca and Doebeli, 2018). As a type of transition matrix in the Mk model, all rate different (ARD) matrix was selected. Second, we identified the branches connecting the parental-state (L) nodes and the mutated-sate (R, Q, or M) nodes (red branches in **Figure S1C**). In these branches, it is expected that the mutation acquisitions in the S L452 residue occurred. Finally, we counted the descendant sequences of respective branches where the mutations in the S L452 were likely acquired. If the number of descendants is ≥10, we defined these descendant sequences as a common ancestry group of the BA.2 variants, which bears a mutation at position 452 in S. Information of the common ancestry group is summarized in **Table S2**.

#### Modeling the epidemic dynamics of SARS-CoV-2 lineages

To quantify the spread rate of each SARS-CoV-2 lineage in the human population, we estimated the relative effective reproduction number of each viral lineage according to the epidemic dynamics, calculated on the basis of viral genomic surveillance data. The data were downloaded from the GISAID database (https://www.gisaid.org/) on May 15, 2022. We excluded the data of viral strains with the following features from the analysis: i) a lack of collection date information; ii) sampling in animals other than humans; or iii) sampling by quarantine. We analyzed the datasets of the five countries (South Africa, the USA, France, Denmark and Belgium) where BA.4/5, BA.2.12.1, BA.2.11, BA.2.9.1, and BA.2.13 were most detected, respectively (**Table S4**). The BA.2 sublineages without amino acid mutations at position 452 in S were summarized as BA.2. In addition, the Delta sublineages were also summarized as Delta. The dynamics of up to five most predominant viral lineages in each country from February 5, 2022, to May 15, 2022, were analyzed. The number of viral sequences of each viral lineage collected on each day in each country was counted, and the count matrix was constructed as an input for the statistical model below.

We constructed a Bayesian statistical model to represent relative lineage growth dynamics with multinomial logistic regression, as described in our previous study (Suzuki *et al*., 2022). In the present study, the epidemic dynamics in respective countries were independently estimated. Arrays in the model index over one or more indices: viral lineages *l* and days *t*. The model is:

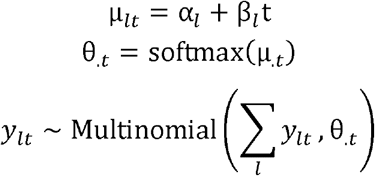

The explanatory variable was time, *t*, and the outcome variable was *y_lt_*, which represented the count of viral lineage *l* at time *t*. In the model, the linear estimator µ_.*t*_, consisting of the intercept α. and the slope β., was converted to the simplex θ_.*t*_, which represented the probability of occurrence of each viral lineage at time *t*, based on the softmax link function defined as:

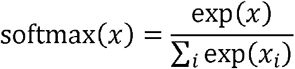

*y_lt_* is generated from θ_.*t*_ and the total count of all lineages at time *t* according to a multinomial distribution.

The relative R_e_ of each viral lineage (*r_l_*) was calculated according to the slope parameter β*_l_* as:

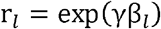

where γ is the average viral generation time (2.1 days) (http://sonorouschocolate.com/covid19/index.php?title=Estimating_Generation_Time_Of_Omicron).

For parameter estimation, the intercept and slope parameters of the BA.2 variant were fixed at 0. Consequently, the relative R_e_ of BA.2 was fixed at 1, and those of the other lineages were estimated relative to that of BA.2.

Parameter estimation was performed via the MCMC approach implemented in CmdStan v2.28.1 (https://mc-stan.org) with CmdStanr v0.4.0 (https://mc-stan.org/cmdstanr/). Noninformative priors were set for all parameters. Four independent MCMC chains were run with 500 and 2,000 steps in the warmup and sampling iterations, respectively. We confirmed that all estimated parameters showed < 1.01 R-hat convergence diagnostic values and > 200 effective sampling size values, indicating that the MCMC runs were successfully convergent. The fitted model closely recapitulated the observed viral lineage dynamics (**Figures 1E and S1D**). The above analyses were performed in R v4.1.3 (https://www.r-project.org/). Information on the relative effective reproduction number of BA.2 estimated in the present study is summarized in **Table S3**.

#### Plasmid construction

Plasmids expressing the SARS-CoV-2 S proteins of B.1.1 (the parental D614G-bearing variant) and BA.2 were prepared in our previous studies (Kimura *et al*., 2022a; Ozono *et al*., 2021; Suzuki *et al*., 2022; Yamasoba *et al*., 2022a). Plasmids expressing the codon-optimized S proteins of L452R/Q/M-bearing variants and their derivatives were generated by site-directed overlap extension PCR using the primers listed in **Table S6**. The resulting PCR fragment was digested with KpnI and NotI and inserted into the corresponding site of the pCAGGS vector (Niwa et al., 1991). A plasmid encoding the SARS-CoV-2 BA.2 S RBD (residues 322-536) was cloned into the expression vector pHLsec containing the N-terminal secretion signal sequence and the C-terminal His^6^-tag sequence (Aricescu et al., 2006). Nucleotide sequences were determined by DNA sequencing services (Eurofins), and the sequence data were analyzed by Sequencher v5.1 software (Gene Codes Corporation).

#### Preparation of BA.2 S RBD

The BA.2 S RBD was prepared as previously described (Kubota *et al*., 2016). Briefly, the expression plasmid encoding the BA.2 S RBD was transfected into 293S GnTI(-) cells (Kubota *et al*., 2016). The proteins in the culture supernatant were purified with cOmplete His-Tag Purification Resin (Roche) affinity column, followed by Superdex 75 Increase 10/300 size-exclusion chromatography (Cytiva) with calcium- and magnesium-free PBS buffer.

#### Preparation of mouse sera

BALB/c mice (female, 7 weeks old) were immunized with 1 μg SARS-CoV-2 BA.2 RBD protein in 50% AddaVax (Invivogen, Cat# vac-adx-10) at day 0 and 14. Ten days after second immunization, blood was collected in BD microtainer blood collection tubes (BD Biosciences, Cat# 365967) and sera were collected by centrifugation.

#### Preparation of human airway and alveolar epithelial cells from human iPSCs

The air-liquid interface culture of airway and alveolar epithelial cells were differentiated from human iPSC-derived lung progenitor cells as previously described (Gotoh et al., 2014; Konishi et al., 2016; Yamamoto et al., 2017). Briefly, lung progenitor cells were stepwise induced from human iPSCs referring a 21-days and 4-steps protocol (Yamamoto *et al*., 2017). At day 21, lung progenitor cells were isolated with specific surface antigen carboxypeptidase M and seeded onto upper chamber of 24-well Cell Culture Insert (Falcon, #353104), followed by 28-day and 7-day differentiation of airway and alveolar epithelial cells, respectively. Alveolar differentiation medium supplemented with dexamethasone (Sigma-Aldrich, Cat# D4902), KGF (PeproTech, Cat# 100-19), 8-Br-cAMP (Biolog, Cat# B007), 3-Isobutyl 1-methylxanthine (IBMX), CHIR99021 (Axon Medchem, Cat# 1386), and SB431542 (FUJIFILM Wako, Cat# 198-16543) was used for induction of alveolar epithelial cells. PneumaCult ALI (STEMCELL Technologies, Cat# ST-05001) supplemented with heparin and Y-27632 (LC Laboratories, Cat# Y-5301) hydrocortisone (Sigma-Aldrich, Cat# H0135) was used for induction of airway epithelial cells.

#### Neutralization assay

Pseudoviruses were prepared as previously described (Kimura *et al*., 2022a; Meng *et al*., 2022; Ozono *et al*., 2021; Saito *et al*., 2022; Uriu et al., 2022; Uriu *et al*., 2021; Yamasoba *et al*., 2022a; Yamasoba *et al*., 2022b). Briefly, lentivirus (HIV-1)-based, luciferase-expressing reporter viruses were pseudotyped with the SARS-CoV-2 S proteins. HEK293T cells (1,000,000 cells) were cotransfected with 1 μg psPAX2-IN/HiBiT (Ozono et al., 2020), 1 μg pWPI-Luc2 (Ozono *et al*., 2020), and 500 ng plasmids expressing parental S or its derivatives using PEI Max (Polysciences, Cat# 24765-1) according to the manufacturer’s protocol. Two days posttransfection, the culture supernatants were harvested and centrifuged. The pseudoviruses were stored at –80°C until use.

Neutralization assay (**Figure 2**) was prepared as previously described (Kimura *et al*., 2022a; Meng *et al*., 2022; Ozono *et al*., 2021; Saito *et al*., 2022; Uriu *et al*., 2022; Uriu *et al*., 2021; Yamasoba *et al*., 2022a; Yamasoba *et al*., 2022b). Briefly, the SARS-CoV-2 S pseudoviruses (counting ∼20,000 relative light units) were incubated with serially diluted (120-fold to 97,480-fold dilution at the final concentration) heat-inactivated sera at 37°C for 1 hour. Pseudoviruses without sera were included as controls. Then, an 40 μl mixture of pseudovirus and serum/antibody was added to HOS-ACE2/TMPRSS2 cells (10,000 cells/50 μl) in a 96-well white plate. At 2 d.p.i., the infected cells were lysed with a One-Glo luciferase assay system (Promega, Cat# E6130) or a Bright-Glo luciferase assay system (Promega, Cat# E2650), and the luminescent signal was measured using a GloMax explorer multimode microplate reader 3500 (Promega) or CentroXS3 (Berthhold Technologies). The assay of each serum was performed in triplicate, and the 50% neutralization titer (NT50) was calculated using Prism 9 software v9.1.1 (GraphPad Software).

#### Pseudovirus infection

Pseudovirus infection was (**Figure 3A**) performed as previously described (Kimura *et al*., 2022a; Kimura *et al*., 2022b; Motozono *et al*., 2021; Saito *et al*., 2022; Suzuki *et al*., 2022; Uriu *et al*., 2022; Uriu *et al*., 2021; Yamasoba *et al*., 2022a; Yamasoba *et al*., 2022b). Briefly, the amount of pseudoviruses prepared was quantified by the HiBiT assay using Nano Glo HiBiT lytic detection system (Promega,Cat# N3040) as previously described (Ozono *et al*., 2021; Ozono *et al*., 2020), and the same amount of pseudoviruses (normalized to the HiBiT value, which indicates the amount of p24 HIV-1 antigen) was inoculated into HOS-ACE2/TMPRSS2 cells, HEK293-ACE2 cells or HEK293-ACE2/TMPRSS2 and viral infectivity was measured as described above (see “Neutralization assay” section). To analyze the effect of TMPRSS2 for pseudovirus infectivity (**Figure S2A**), the fold change of the values of HEK293-ACE2/TMPRSS2 to HEK293-ACE2 was calculated.

#### Yeast surface display

Yeast surface display (**Figure 3B**) was performed as previously described as previously described (Dejnirattisai *et al*., 2022; Kimura *et al*., 2022a; Kimura *et al*., 2022b; Motozono *et al*., 2021; Yamasoba *et al*., 2022a; Zahradnik *et al*., 2021a). Briefly, yeast codon-optimized SARS-CoV-2_RBD-Omicron-BA.2 was obtained from Twist Biosciences and the mutant RBDs were PCR amplified by KAPA HiFi HotStart ReadyMix kit (Roche, Cat# KK2601) and assembled in vivo by yeast [*Saccharomyces cerevisiae* strain EBY100 (ATCC, MYA-4941)] homologous recombination with pJYDC1 plasmid (Addgene, Cat# 162458) as previously described (Dejnirattisai *et al*., 2022; Kimura *et al*., 2022a; Kimura *et al*., 2022b; Motozono *et al*., 2021; Yamasoba *et al*., 2022a; Zahradnik *et al*., 2021a). Primers used are listed in **Table S6**. Yeasts were expressed for 48 hours at 20°C, washed with PBS supplemented with bovine serum albumin at 1 g/l and incubated with 12–14 different concentrations of Expi293F cells produced ACE2 peptidase domain (residues 18-740, 200 nM to 13 pM) for 12 hours. To induce eUnaG2 reporter protein fluorescence, bilirubin (Sigma-Aldrich, Cat# 14370-1G) was added to the final concentration of 5 nM. RBD expression and ACE2 signal were recorded by using a FACS S3e cell sorter device (Bio-Rad), background binding signals were subtracted and data were fitted to a standard noncooperative Hill equation by nonlinear least-squares regression using Python v3.7 (https://www.python.org) as previously described (Kimura *et al*., 2022a; Kimura *et al*., 2022b; Motozono *et al*., 2021; Yamasoba *et al*., 2022a; Zahradnik et al., 2021b).

#### SARS-CoV-2 S-based fusion assay

SARS-CoV-2 S-based fusion assay (**Figures 3D and S2C**) was performed as previously described (Kimura *et al*., 2022b; Motozono *et al*., 2021; Saito *et al*., 2022; Suzuki *et al*., 2022; Yamasoba *et al*., 2022a). Briefly, on day 1, effector cells (i.e., S-expressing cells) and target cells (see below) were prepared at a density of 0.6–0.8 × 10^6^ cells in a 6-well plate. To prepare effector cells, HEK293 cells were cotransfected with the S expression plasmids (400 ng) and pDSP_8-11_ (Kondo et al., 2011) (400 ng) using TransIT-LT1 (Takara, Cat# MIR2300). On day 2, to prepare target cells, VeroE6/TMPRSS2 cells were transfected with pDSP_1-7_ (Kondo *et al*., 2011) (400 ng). On day 3 (24 hours posttransfection), 16,000 effector cells were detached and reseeded into 96-well black plates (PerkinElmer, Cat# 6005225), and target cells (VeroE6/TMPRSS2 or Calu-3/DSP_1-7_ cells) were reseeded at a density of 1,000,000 cells/2 ml/well in 6-well plates. On day 4 (48 hours posttransfection), target cells were incubated with EnduRen live cell substrate (Promega, Cat# E6481) for 3 hours and then detached, and 32,000 target cells were added to a 96-well plate with effector cells. *Renilla* luciferase activity was measured at the indicated time points using Centro XS3 LB960 (Berthhold Technologies). To measure the surface expression level of S protein, effector cells were stained with rabbit anti-SARS-CoV-2 S S1/S2 polyclonal antibody (Thermo Fisher Scientific, Cat# PA5-112048, 1:100). Normal rabbit IgG (SouthernBiotech, Cat# 0111-01, 1:100) was used as negative controls, and APC-conjugated goat anti-rabbit IgG polyclonal antibody (Jackson ImmunoResearch, Cat# 111-136-144, 1:50) was used as a secondary antibody. Surface expression level of S proteins (**Figures 3C and S2B**) was measured using FACS Canto II (BD Biosciences) and the data were analyzed using FlowJo software v10.7.1 (BD Biosciences). To calculate fusion activity, *Renilla* luciferase activity was normalized to the MFI of surface S proteins. The normalized value (i.e., *Renilla* luciferase activity per the surface S MFI) is shown as fusion activity.

#### Coculture experiment

Coculture experiment (**Figure S2D**) was performed as previously described (Suzuki *et al*., 2022; Yamasoba *et al*., 2022a). This assay utilizes a dual split protein (DSP) encoding *Renilla* luciferase and *GFP* genes; the respective split proteins, DSP_8-11_ and DSP_1-7_, are expressed in effector and target cells by transfection. Briefly, one day before transfection, effector cells (i.e., S-expressing cells) were seeded on the poly-L-lysine (Sigma, Cat# P4832) coated coverslips put in a 12-well plate, and target cells were prepared at a density of 100,000 cells in a 12-well plate. To prepare effector cells, HEK293 cells were cotransfected with the S-expression plasmids (500 ng) and pDSP_8-11_ (500 ng) using PEI Max (Polysciences, Cat# 24765-1). To prepare target cells, HEK293 and HEK293-ACE2/TMPRSS2 cells were transfected with pDSP_1-7_ (500 ng) (Kondo *et al*., 2011). At 24 hours posttransfection, target cells were detached and cocultured with effector cells in a 1:2 ratio. At 9 h post-coculture, cells were fixed with 4% paraformaldehyde in PBS (Nacalai Tesque, Cat# 09154-85) for 15 minutes at room temperature. Nuclei were stained with Hoechst 33342 (Thermo Fisher Scientific, Cat# H3570). The coverslips were mounted on glass slides using Fluoromount-G (SouthernBiotech, Cat# 0100-01) with Hoechst 33342 and observed using an A1Rsi Confocal Microscope (Nikon). The size of syncytium (GFP-positive area) was measured using Fiji software v2.2.0 (ImageJ) as previously described (Suzuki *et al*., 2022; Yamasoba *et al*., 2022a).

#### SARS-CoV-2 reverse genetics

Recombinant SARS-CoV-2 was generated by circular polymerase extension reaction (CPER) as previously described (Kimura *et al*., 2022b; Motozono *et al*., 2021; Saito *et al*., 2022; Torii *et al*., 2021; Yamasoba *et al*., 2022a). To generate the lineage A-based GFP-expressing chimeric recombinant SARS-CoV-2 (rBA.2.9.1 S-GFP, rBA.2.11 S-GFP, rBA.2.12.1 S-GFP and rBA.4/5 S-GFP) (**Figure 4A, left**), 7 DNA fragments (fragments 1-7) encoding the partial genome of SARS-CoV-2 (strain WK-521, PANGO lineage A; GISAID ID: EPI_ISL_408667) (Matsuyama *et al*., 2020) were prepared by PCR using PrimeSTAR GXL DNA polymerase (Takara, Cat# R050A). A linker fragment encoding hepatitis delta virus ribozyme, bovine growth hormone poly A signal and cytomegalovirus promoter was also prepared by PCR. The corresponding SARS-CoV-2 genomic region and the PCR templates and primers used for this procedure are summarized in **Table S6**. Instead of the authentic fragment 9, we used the fragment 9, in which the *GFP* gene was inserted in the *ORF7a* frame (**Table S6**) (Torii *et al*., 2021). To prepare the fragment 8 encoding the *S* genes of BA.2.9.1, BA.2.11, BA.2.12.1 and BA.4/5, we inserted additional mutations into the fragment 8 plasmid encoding the BA.2 S (Yamasoba *et al*., 2022a) by site-directed overlap extension PCR using the primers listed in **Table S6**. Nucleotide sequences were determined by a DNA sequencing service (Fasmac), and the sequence data were analyzed by Sequencher software v5.1 (Gene Codes Corporation). Finally, the 10 DNA fragments were mixed and used for CPER (Torii *et al*., 2021).

To generate the BA.2-based chimeric recombinant SARS-CoV-2 (rBA.2, rBA.2.9.1, rBA.2.11 and rBA.4/5) (**Figure 4A, right**), RNA was extracted from the cells infected with a clinical isolate of BA.2 (GISAID ID: EPI_ISL_9595859) and cDNA was synthesized as described above (see “Viral genome sequencing” section). The two DNA fragments correspond to the fragments 1-7 and 9 were prepared by RT-PCR using PrimeSTAR GXL DNA polymerase (Takara, Cat# R050A) using the primers listed in **Table S6**. The fragments 8 bearing the *S* genes of BA.2, BA.2.9.1, BA.2.11, BA.2.12.1 and BA.4/5 were prepared as described above. Finally, the 3 DNA fragments were mixed and used for CPER (Torii *et al*., 2021).

To produce recombinant SARS-CoV-2 (seed viruses), the CPER products were transfected into HEK293-C34 cells using TransIT-LT1 (Takara, Cat# MIR2300) according to the manufacturer’s protocol. At one day posttransfection, the culture medium was replaced with DMEM (high glucose) (Sigma-Aldrich, Cat# 6429-500ML) containing 2% FBS, 1% PS and doxycycline (1 μg/ml; Takara, Cat# 1311N). At six days posttransfection, the culture medium was harvested and centrifuged, and the supernatants were collected as the seed virus. To remove the CPER products (i.e., SARS-CoV-2-related DNA), 1 ml of the seed virus was treated with 2 μl TURBO DNase (Thermo Fisher Scientific, Cat# AM2238) and incubated at 37°C for 1 hour. Complete removal of the CPER products from the seed virus was verified by PCR. The working virus stock was prepared using the seed virus as described below (see “SARS-CoV-2 preparation and titration” section).

#### SARS-CoV-2 preparation and titration

The working virus stocks of chimeric recombinant SARS-CoV-2 were prepared and titrated as previously described (Kimura *et al*., 2022b; Motozono *et al*., 2021; Saito *et al*., 2022; Torii *et al*., 2021; Yamasoba *et al*., 2022a). In brief, 20 μl of the seed virus was inoculated into VeroE6/TMPRSS2 cells (5,000,000 cells in a T-75 flask). One hour postinfection (h.p.i.), the culture medium was replaced with DMEM (low glucose) (Wako, Cat# 041-29775) containing 2% FBS and 1% PS. At 3 d.p.i., the culture medium was harvested and centrifuged, and the supernatants were collected as the working virus stock.

The titer of the prepared working virus was measured as the 50% tissue culture infectious dose (TCID_50_). Briefly, one day before infection, VeroE6/TMPRSS2 cells (10,000 cells) were seeded into a 96-well plate. Serially diluted virus stocks were inoculated into the cells and incubated at 37°C for 4 days. The cells were observed under microscopy to judge the CPE appearance. The value of TCID_50_/ml was calculated with the Reed–Muench method (Reed and Muench, 1938).

To verify the sequences of SARS-CoV-2 working viruses, viral RNA was extracted from the working viruses using a QIAamp viral RNA mini kit (Qiagen, Cat# 52906) and viral genome sequences were analyzed as described above (see “Viral genome sequencing” section). Information on the unexpected mutations detected is summarized in **Table S7**, and the raw data are deposited in the GitHub repository (https://github.com/TheSatoLab/BA.2_related_Omicrons).

#### Plaque assay

Plaque assay (**Figures 4B and 4C**) was performed as previously described (Kimura *et al*., 2022b; Motozono *et al*., 2021; Saito *et al*., 2022; Suzuki *et al*., 2022; Yamasoba *et al*., 2022a). Briefly, one day before infection, VeroE6/TMPRSS2 cells (100,000 cells) were seeded into a 24-well plate and infected with SARS-CoV-2 (1, 10, 100 and 1,000 TCID_50_) at 37°C for 1 hour. Mounting solution containing 3% FBS and 1.5% carboxymethyl cellulose (Wako, Cat# 039-01335) was overlaid, followed by incubation at 37°C. At 3 d.p.i., the culture medium was removed, and the cells were washed with PBS three times and fixed with 4% paraformaldehyde phosphate (Nacalai Tesque, Cat# 09154-85). The fixed cells were washed with tap water, dried, and stained with staining solution [0.1% methylene blue (Nacalai Tesque, Cat# 22412-14) in water] for 30 minutes. The stained cells were washed with tap water and dried, and the size of plaques was measured using Fiji software v2.2.0 (ImageJ).

#### SARS-CoV-2 infection

One day before infection, Vero cells (10,000 cells) and VeroE6/TMPRSS2 cells (10,000 cells) were seeded into a 96-well plate. SARS-CoV-2 [1,000 TCID_50_ for Vero cells (**Figures 4D and 4F**); 100 TCID_50_ for VeroE6/TMPRSS2 cells (**Figures 4E and 4G**)] was inoculated and incubated at 37°C for 1 hour. The infected cells were washed, and 180 µl culture medium was added. The culture supernatant (10 µl) was harvested at the indicated timepoints and used for RT–qPCR to quantify the viral RNA copy number (see “RT–qPCR” section below) In the infection experiment using human iPSC-derived airway and alveolar epithelial cells (**Figures 4H and 4I**), working viruses were diluted with Opti-MEM (Thermo Fisher Scientific, 11058021). The diluted viruses (1,000 TCID_50_ in 100□μl) were inoculated onto the apical side of the culture and incubated at 37□°C for 1□hour. The inoculated viruses were removed and washed twice with Opti-MEM. To collect the viruses, 100□μl Opti-MEM was applied onto the apical side of the culture and incubated at 37□°C for 10□minutes. The Opti-MEM was collected and used for RT–qPCR to quantify the viral RNA copy number (see “RT–qPCR” section below).

#### RT–qPCR

RT–qPCR was performed as previously described (Kimura *et al*., 2022b; Meng *et al*., 2022; Motozono *et al*., 2021; Saito *et al*., 2022; Suzuki *et al*., 2022; Yamasoba *et al*., 2022a). Briefly, 5 μl culture supernatant was mixed with 5 μl 2 × RNA lysis buffer [2% Triton X-100 (Nacalai Tesque, Cat# 35501-15), 50 mM KCl, 100 mM Tris-HCl (pH 7.4), 40% glycerol, 0.8 U/μl recombinant RNase inhibitor (Takara, Cat# 2313B)] and incubated at room temperature for 10 m. RNase-free water (90 μl) was added, and the diluted sample (2.5 μl) was used as the template for real-time RT-PCR performed according to the manufacturer’s protocol using One Step TB Green PrimeScript PLUS RT-PCR kit (Takara, Cat# RR096A) and the following primers: Forward *N*, 5’-AGC CTC TTC TCG TTC CTC ATC AC-3’; and Reverse *N*, 5’-CCG CCA TTG CCA GCC ATT C-3’. The viral RNA copy number was standardized with a SARS-CoV-2 direct detection RT-qPCR kit (Takara, Cat# RC300A). Fluorescent signals were acquired using QuantStudio 3 Real-Time PCR system (Thermo Fisher Scientific), CFX Connect Real-Time PCR Detection system (Bio-Rad), Eco Real-Time PCR System (Illumina), qTOWER3 G Real-Time System (Analytik Jena) or 7500 Real-Time PCR System (Thermo Fisher Scientific).

#### Animal experiments

Animal experiments (**Figure 5**) were performed as previously described (Saito *et al*., 2022; Suzuki *et al*., 2022; Yamasoba *et al*., 2022a). Syrian hamsters (male, 4 weeks old) were purchased from Japan SLC Inc. (Shizuoka, Japan). Baseline body weights were measured before infection. For the virus infection experiments, hamsters were anaesthetized by intramuscular injection of a mixture of either 0.15 mg/kg medetomidine hydrochloride (Domitor^®^, Nippon Zenyaku Kogyo), 2.0 mg/kg midazolam (FUJIFILM Wako Chemicals, Cat# 135-13791) and 2.5 mg/kg butorphanol (Vetorphale^®^, Meiji Seika Pharma), or 0.15 mg/kg medetomidine hydrochloride, 2.0 mg/kg alphaxaone (Alfaxan^®^, Jurox) and 2.5 mg/kg butorphanol. The chimeric recombinant SARS-CoV-2 (rBA.2, rBA.2.12.1, and rBA.4/5) (10,000 TCID_50_ in 100 µl), or saline (100 µl) were intranasally inoculated under anesthesia. Oral swabs were collected at 1, 3, and 5 d.p.i. Oral swabs were daily collected under anesthesia with isoflurane (Sumitomo Dainippon Pharma). Body weight, enhanced pause (Penh), the ratio of time to peak expiratory follow relative to the total expiratory time (Rpef) and subcutaneous oxygen saturation (SpO_2_) were routinely monitored at indicated timepoints (see “Lung function test” section below). Respiratory organs were anatomically collected at 1, 3 and 5 d.p.i (for lung) or 1 d.p.i. (for trachea). Viral RNA load in the respiratory tissues and oral swab were determined by RT–qPCR. The respiratory tissues were also used for histopathological and IHC analyses (see “H&E staining” and “IHC” sections below). Sera of infected hamsters were collected at 16 d.p.i. using cardiac puncture under anesthesia with isoflurane and used for neutralization assay (see “Neutralization assay” above).

#### Lung function test

Lung function test (**Figure 5A**) was performed at 1, 3, 5, and 7 d.p.i. as previously described (Suzuki *et al*., 2022; Yamasoba *et al*., 2022a). Respiratory parameters (Penh and Rpef) were measured by using a whole-body plethysmography system (DSI) according to the manufacturer’s instructions. In brief, a hamster was placed in an unrestrained plethysmography chamber and allowed to acclimatize for 30 seconds, then, data were acquired over a 2.5-minute period by using FinePointe Station and Review softwares v2.9.2.12849 (STARR). The state of oxygenation was examined by measuring SpO_2_ using pulse oximeter, MouseOx PLUS (STARR). SpO_2_ was measured by attaching a measuring chip to the neck of hamsters sedated by 0.25 mg/kg medetomidine hydrochloride.

#### IHC

IHC (**Figures 5D, S3A and S3B**) was performed as previously described (Saito *et al*., 2022; Suzuki *et al*., 2022; Yamasoba *et al*., 2022a) using an Autostainer Link 48 (Dako). The deparaffinized sections were exposed to EnVision FLEX target retrieval solution high pH (Agilent, Cat# K8004) for 20 minutes at 97°C to activate, and mouse anti-SARS-CoV-2 N monoclonal antibody (clone 1035111, R&D systems, Cat# MAB10474-SP, 1:400) was used as a primary antibody. The sections were sensitized using EnVision FLEX (Agilent) for 15 minutes and visualized by peroxidase-based enzymatic reaction with 3,3’-diaminobenzidine tetrahydrochloride (Dako, Cat# DM827) as substrate for 5 minutes. The N protein positivity (**Figures 5E and S3B**) was evaluated by certificated pathologists as previously described (Suzuki *et al*., 2022; Yamasoba *et al*., 2022a). Images were incorporated as virtual slide by NDP.scan software v3.2.4 (Hamamatsu Photonics). The N-protein positivity was measured as the area using Fiji software v2.2.0 (ImageJ).

#### H&E staining

H&E staining (**Figure 5G**) was performed as previously described (Saito *et al*., 2022; Suzuki *et al*., 2022; Yamasoba *et al*., 2022a). Briefly, excised animal tissues were fixed with 10% formalin neutral buffer solution, and processed for paraffin embedding. The paraffin blocks were sectioned with 3 µm-thickness and then mounted on MAS-GP-coated glass slides (Matsunami Glass, Cat# S9901). H&E staining was performed according to a standard protocol.

#### Histopathological scoring

Histopathological scoring (**Figure 5F**) was performed as previously described (Saito *et al*., 2022; Suzuki *et al*., 2022; Yamasoba *et al*., 2022a). Pathological features including bronchitis or bronchiolitis, hemorrhage with congestive edema, alveolar damage with epithelial apoptosis and macrophage infiltration, hyperplasia of type II pneumocytes, and the area of the hyperplasia of large type II pneumocytes were evaluated by certified pathologists and the degree of these pathological findings were arbitrarily scored using four-tiered system as 0 (negative), 1 (weak), 2 (moderate), and 3 (severe). The “large type II pneumocytes” are the hyperplasia of type II pneumocytes exhibiting more than 10-μm-diameter nucleus. We described “large type II pneumocytes” as one of the remarkable histopathological features reacting SARS-CoV-2 infection in our previous studies (Saito *et al*., 2022; Suzuki *et al*., 2022; Yamasoba *et al*., 2022a). Total histology score is the sum of these five indices.

To measure the inflammation area in the infected lungs (**Figures 5H and S3C**), four hamsters infected with each virus were sacrificed at 5 d.p.i., and all four right lung lobes, including upper (anterior/cranial), middle, lower (posterior/caudal), and accessory lobes, were sectioned along with their bronchi. The tissue sections were stained by H&E, and the digital microscopic images were incorporated into virtual slides using NDP.scan software v3.2.4 (Hamamatsu Photonics). The inflammatory area including type II pneumocyte hyperplasia in the infected whole lungs was morphometrically analyzed using Fiji software v2.2.0 (ImageJ).

### QUANTIFICATION AND STATISTICAL ANALYSIS

Statistical significance was tested using a two-sided Mann–Whitney *U*-test, a two-sided Student’s *t*-test or a two-sided paired *t-*test unless otherwise noted. The tests above were performed using Prism 9 software v9.1.1 (GraphPad Software).

In the time-course experiments (**Figures 3D, 4D–4I, 5A–5C, 5F, and S2C**), a multiple regression analysis including experimental conditions (i.e., the types of infected viruses) as explanatory variables and timepoints as qualitative control variables was performed to evaluate the difference between experimental conditions thorough all timepoints. The initial time point was removed from the analysis. *P* value was calculated by a two-sided Wald test. Subsequently, familywise error rates (FWERs) were calculated by the Holm method. These analyses were performed in R v4.1.2 (https://www.r-project.org/).

In **Figures 5D, 5G and S3**, photographs shown are the representative areas of at least two independent experiments by using four hamsters at each timepoint. In **Figure S3A**, photographs shown are the representatives of >20 fields of view taken for each sample.

