## Supplementary figures and images for "Virological characteristics of the novel SARS-CoV-2 Omicron variants including BA.2.12.1, BA.4 and BA.5"

### Figure S1

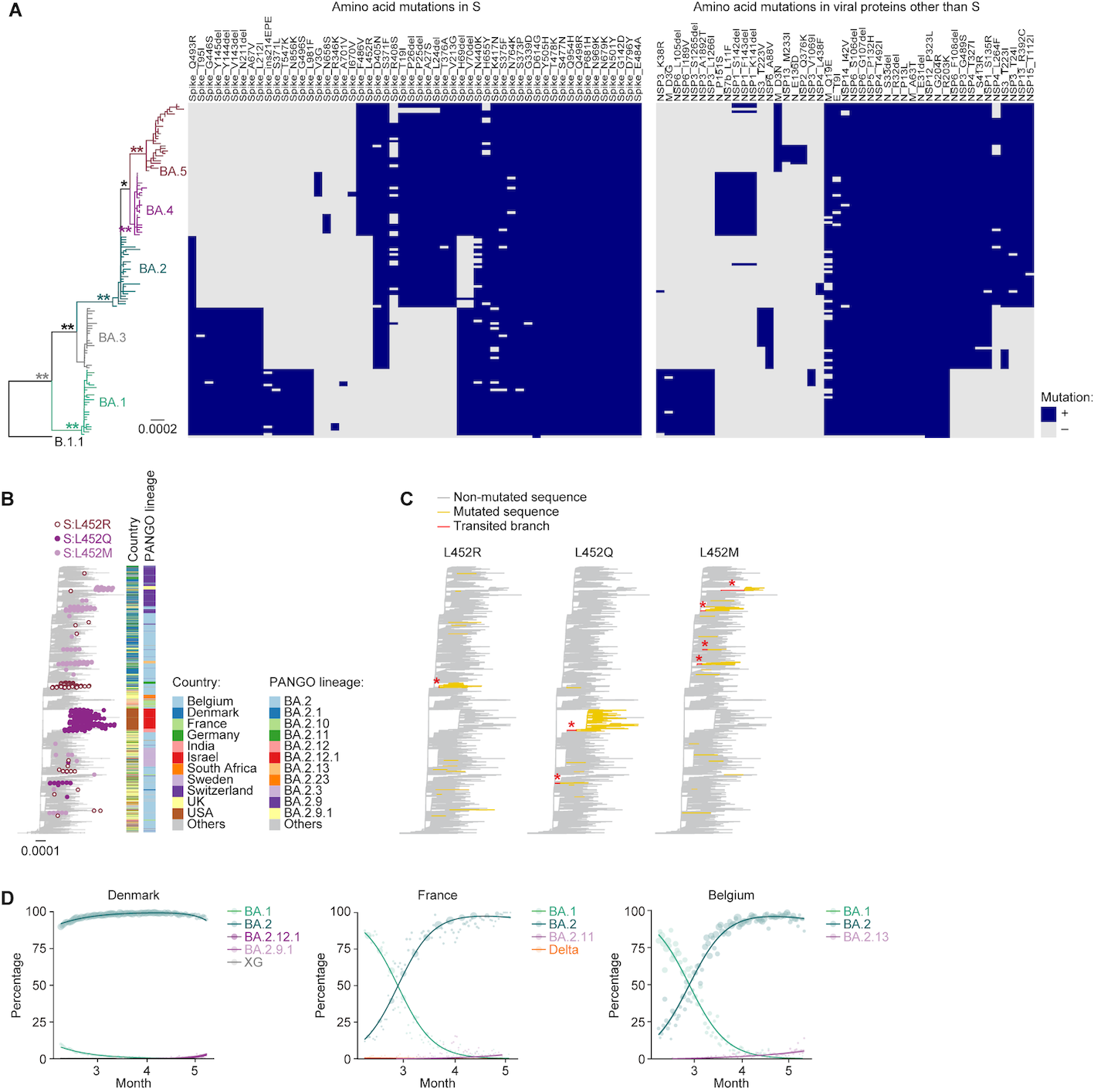

### Figure S2

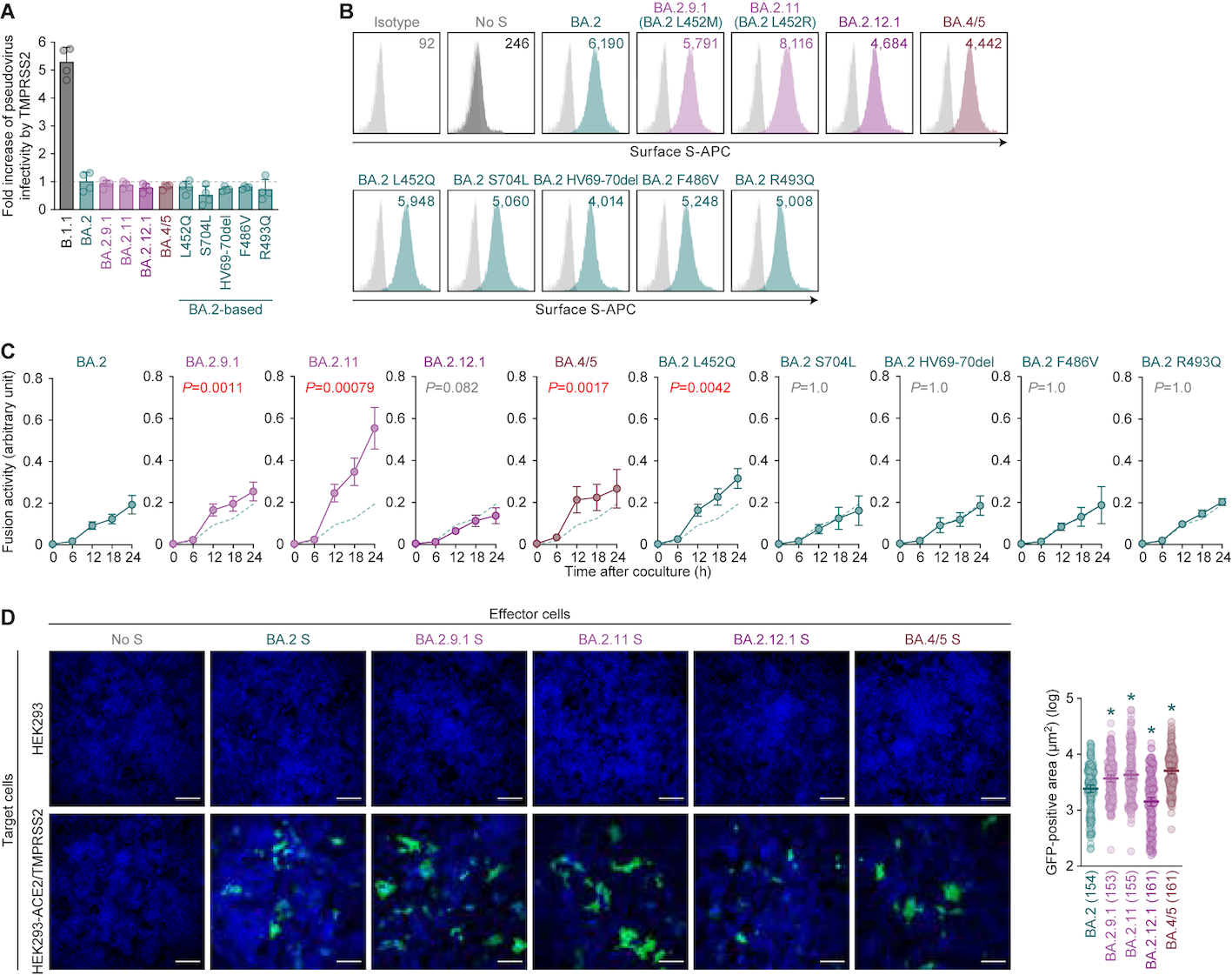

### Figure S3

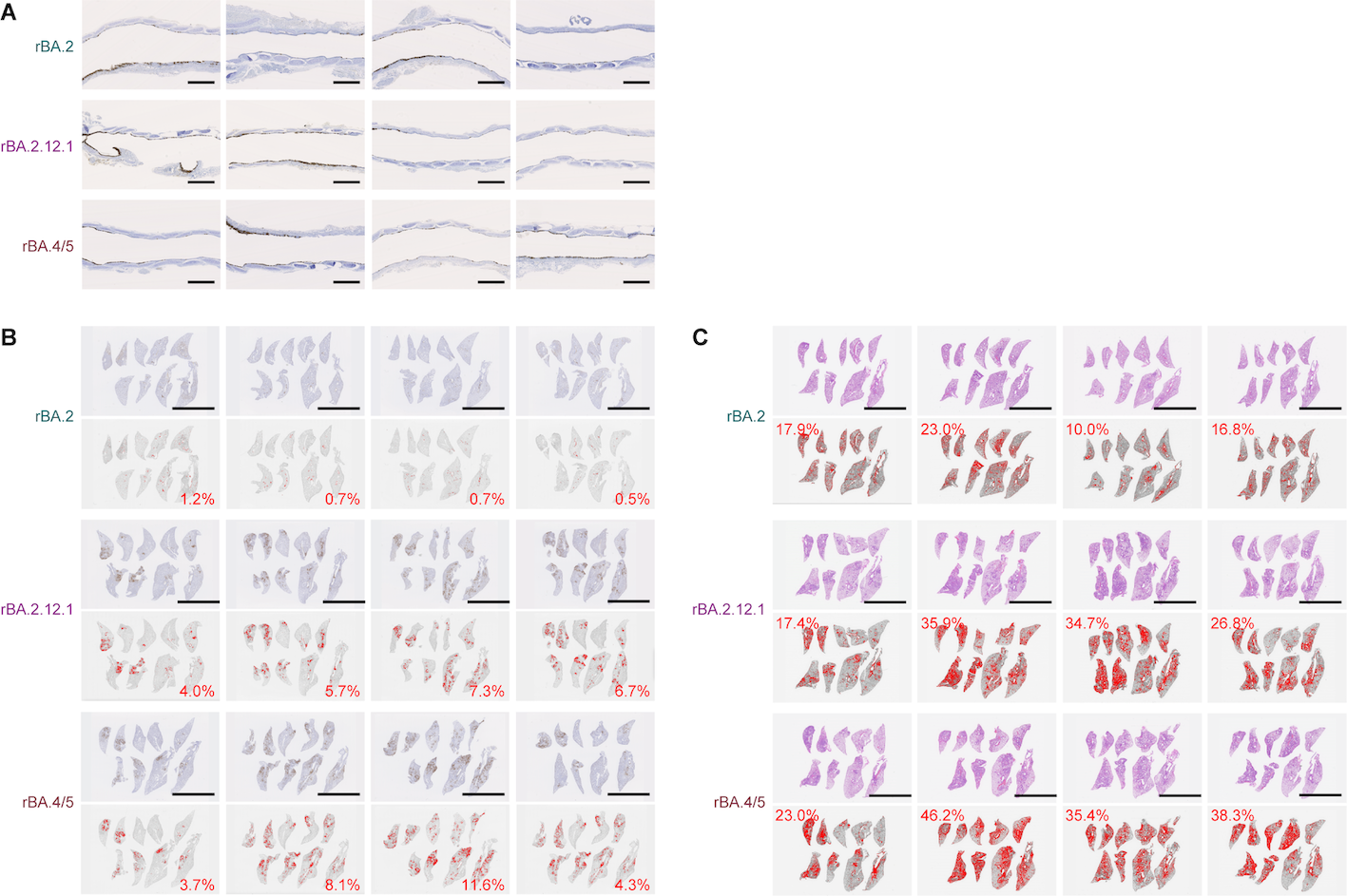
